# Design, synthesis, and pharmacological evaluation of novel PROTAC degraders targeting 11β-HSD1 for metabolic disease intervention

**DOI:** 10.1101/2025.11.09.687522

**Authors:** Liguo Wang, Xue Tao, Ming He, Yu Lu, Yuji Wang, Juanjuan Zhu

## Abstract

Hydroxysteroid 11-beta dehydrogenase 1 (11β-HSD1) plays a critical role in metabolic homeostasis by catalyzing the intracellular conversion of cortisone to cortisol. Dysregulated 11β-HSD1 activity is closely associated with metabolic disorders such as type 2 diabetes mellitus, obesity, and glucocorticoid-related inflammation. While small-molecule inhibitors of 11β-HSD1 have shown promise, they primarily suppress enzymatic activity without modulating protein abundance. Here, we report the development of the 11β-HSD1-targeting PROTAC degraders. A series of bifunctional molecules were synthesized based on CRBN- and VHL-recruiting ligands, with AZD8329-derived warheads linked *via* polyethylene glycol chains. Cellular assays demonstrated efficient, ubiquitin-proteasome-dependent degradation of 11β-HSD1, with H-3-V identified as the most potent degrader. *In vivo*, H-3-V treatment improved glucose tolerance and enhanced glucose-stimulated insulin secretion in a high-fat diet-induced T2DM mouse model. Molecular dynamics simulations revealed that the H-3-V ternary complex exhibited superior binding energetics compared to less active analogs. Collectively, this study introduces a novel chemical modality for 11β-HSD1 modulation and lays the groundwork for future therapeutic development targeting metabolic disease *via* selective protein degradation.

**Graphical abstract:** 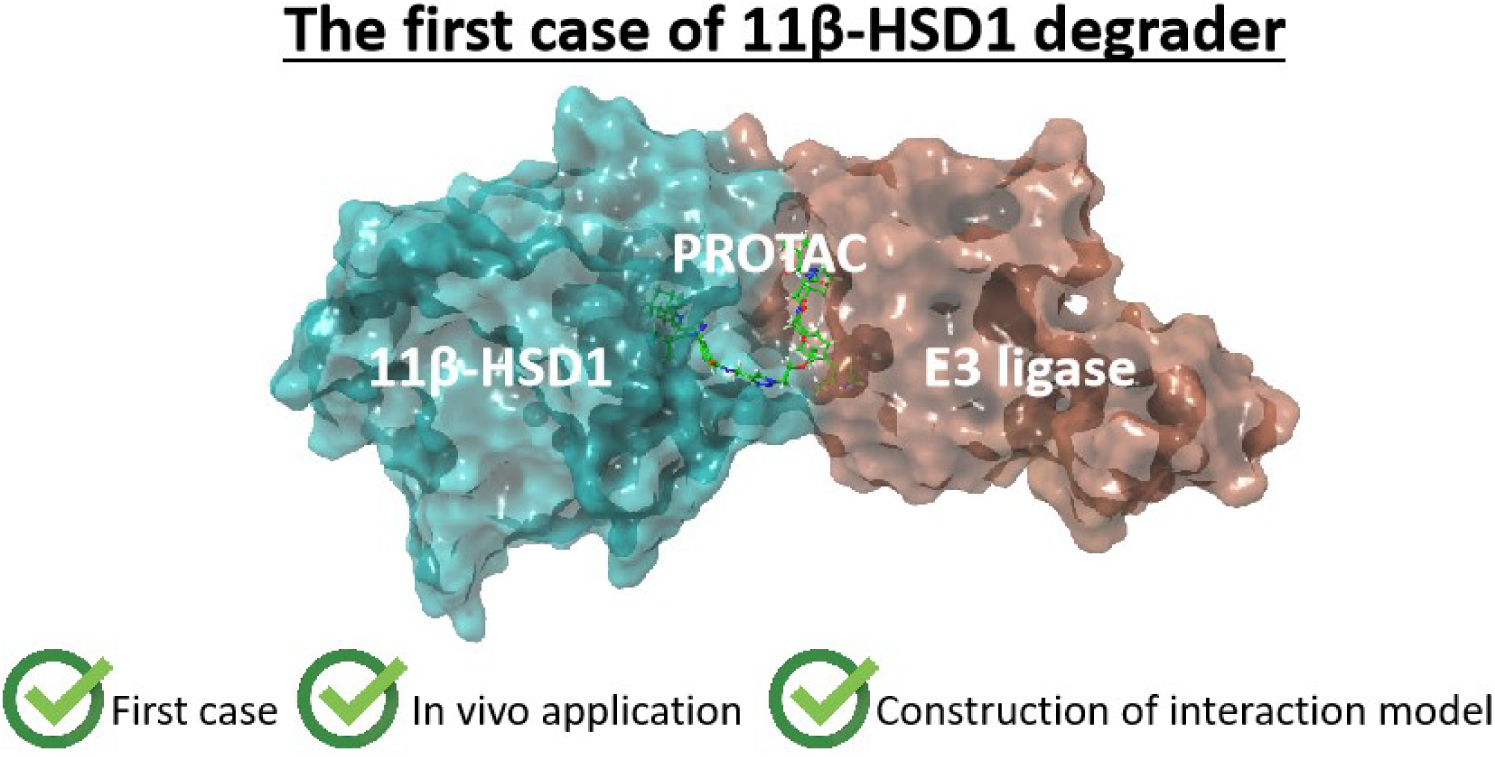

## 1. Introduction

Hydroxysteroid 11-beta dehydrogenase 1 (11β-HSD1), a member of the short-chain dehydrogenase/reductase superfamily, functions as an NADPH-dependent enzyme exhibiting cortisone reductase activity[1]. This enzyme catalyzes the conversion of inactive cortisone to its biologically active reduction product, cortisol. 11β-HSD1 is abundantly expressed in key metabolic tissues, thereby amplifying glucocorticoid action within specific cellular environments[2, 3]. Notably, 11β-HSD1 mediates a bidirectional reaction, maintaining a critical equilibrium between cortisol and cortisone concentrations[4].

Cortisol serves as the primary glucocorticoid (GC), a steroid hormone secreted by the adrenal cortex[5]. For a long time, it has been recognized as playing an indispensable role in regulating carbohydrate, lipid, and protein metabolism, while upholding homeostasis under normal physiological conditions. Critically, overexpression of 11β-HSD1 is strongly associated with glucocorticoid excess, manifesting clinically as Cushing’s syndrome[6]. In experimental models of diabetes, elevated 11β-HSD1 activity exacerbates metabolic dysfunction through impairment of insulin secretion and pancreatic β-cell function[7]. Furthermore, Masuzaki et al. demonstrated that adipose tissue-specific overexpression of 11β-HSD1 induces visceral obesity, insulin resistance, diabetes, dyslipidemia, and hypertension in murine models[8]. Conversely, genetic disruption of 11β-HSD1 (11β-HSD1L/L) confers enhanced glucose tolerance, attenuated hepatic gluconeogenesis, and improved lipid profiles[9, 10].

Moreover, 11β-HSD1 significantly influences the pathogenesis of inflammation-related diseases by modulating intracellular glucocorticoid levels, which critically regulate inflammatory responses[11]. Its overexpression is frequently implicated in chronic inflammatory states, contributing to conditions including obesity-associated inflammation, insulin resistance, atherosclerosis, and metabolic syndrome. In contrast, pharmacological inhibition or genetic deletion of 11β-HSD1 exerts anti-inflammatory effects, enhancing glucose tolerance, reducing tissue-specific inflammation, and attenuating the progression of disorders such as diabetes and cardiovascular disease[12]. Given the pivotal physiological and metabolic functions of 11β-HSD1, numerous pharmaceutical entities are actively developing selective inhibitors targeting this enzyme (**Figure 1A**)[13–17], with several molecular entities advancing into clinical trials.

**Figure 1.**
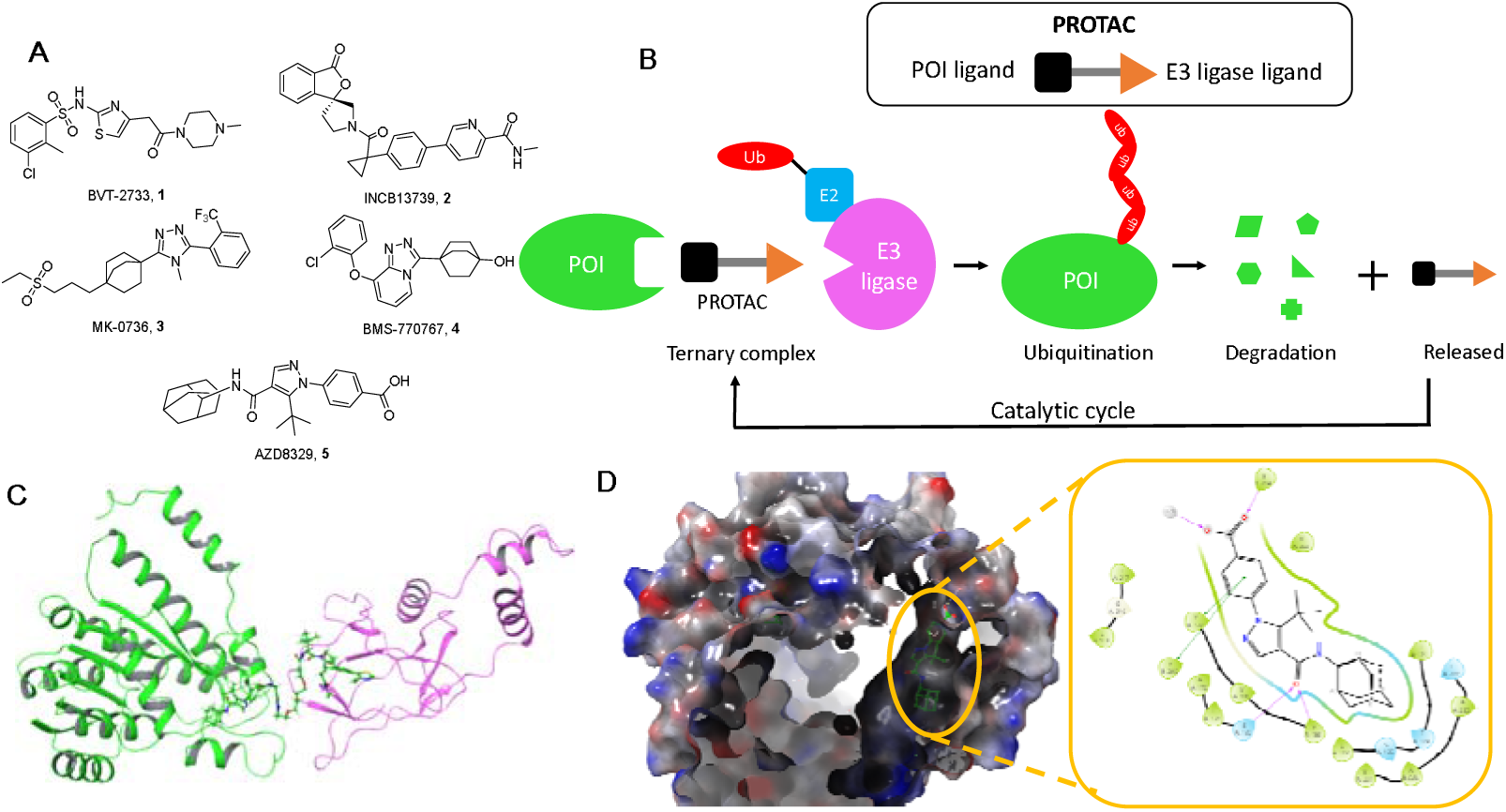
Molecular structure information and PROTAC mechanism of action. (A) Representative molecular structure of 11β-HSD1 inhibitors; (B) Principle of PROTAC molecule action; (C) Schematic diagram of the structure of PROTAC molecules targeting 11β-HSD1. Green: 11β-HSD1 (PDB code: 4P38), pink: von Hippel-Lindau protein (PDB code: 8BDI); (D) Interaction relationship between AZD8329 and 11β-HSD1 (PDB code: 4P38).

These findings identify 11β-HSD1 as a promising therapeutic target for the intervention of metabolic disorders. Recent studies conducted by multiple academic institutions and pharmaceutical companies have provided new insights into 11β-HSD1. Emerging evidence suggests that 11β-HSD1 may have functions beyond its enzymatic activity, particularly through protein - protein interactions and the modulation of cellular signaling pathways. For instance, research shows that 11β-HSD1 interacts with H6PDH proteins, regulating the catalytic oxidation - reduction equilibrium of 11β-HSD[18]. Additionally, Fan Yao et al. demonstrated that H6PDH and 11β-HSD1 are jointly involved in the pathogenesis of type 2 diabetes mellitus and may serve as promising targets for metabolic syndromes[19]. Although significant progress has been made in the development of 11β-HSD1 inhibitors, there is currently no pharmacological agent available to modulate the protein abundance of 11β-HSD1; instead, existing agents directly interfere with the enzymatic function of 11β-HSD1.

Proteolysis-targeting chimera (PROTAC) technology represents an emerging strategy utilizing bifunctional small molecules to achieve targeted protein degradation[20]. A PROTAC molecule comprises three structural elements: a protein-of-interest (POI) ligand, an E3 ubiquitin ligase ligand, and a linker (Figure 1B). This molecular configuration facilitates the formation of a ternary complex between the POI and E3 ligase, ultimately inducing POI ubiquitination and degradation *via* the ubiquitin-proteasome system. To date, several E3 ligase ligands—including those targeting CRBN, VHL, MDM2, cIAP1, and DCAF15—have been successfully implemented in PROTAC development[20]. Herein, we report the first 11β-HSD1 degrader and its successful application in metabolic disorders using PROTAC technology.

## 2. Results and discussion

### 2.1 Molecular design

Given that PROTACs facilitate the interaction between the protein of interest (POI) and the E3 ubiquitin ligase, the identification of suitable ligands, appropriate attachment positions, and optimally designed linkers is essential. As illustrated in **Figure 1C**, several known 11β-HSD1 inhibitors were evaluated as potential ligands for targeting 11β-HSD1. AZD8329 was selected based on its previously characterized binding mode, as reported by Ogg et al. (**Figure 1D**)[21]. The alkyne attachment point was incorporated *via* amide condensation at the protein surface-exposed position of the ligand. In particular, we compared the structural contributions of 1-adamantyl and 2-adamantyl, with XLogP[22] surfaces (1-adamantyl 2.4 *vs.* 2-adamantyl 1.7), and found that 1-adamantyl may possess more hydrophilic characteristics, thereby generating stronger interactions with the hydrophobic regions of amino acids. In molecular design and simulation, substituting 1-adamantyl (QPPCaco[23] 85.6) is likely to produce more favourable cell permeability than 2-adamantyl (QPPCaco 79.7) did in molecular calculation, while this modification did not produce significant conformational differences in binding pattern during molecular simulation (**Supplementary Figure S1**). Therefore, we replaced the 2-adamantyl structure in the AZD8329 structure with 1-adamantyl.

### 2.2 Chemistry

Generally, the degradation activity of PROTAC molecules exhibits considerable variation depending on the selected E3 ubiquitin ligase type. In this study, we employed two distinct classes of PROTAC ligands: pomalidomide-based ligands recruiting CRBN and VH032-based ligands recruiting VHL. Subsequently, we synthesized a series of universal E3 ubiquitin ligase ligands featuring varying linker lengths and terminal azide functionalities (**Figure 2**). These ligands were then conjugated to the protein-of-interest (POI) ligand *via* copper-catalyzed azide-alkyne cycloaddition ("click" reaction), thereby assembling degraders incorporating either VHL or CRBN ligands with differing linker lengths (**Scheme 1**). The synthetic route is shown in **Scheme 2** and **Scheme 3**. Specifically, pomalidomide served as the CRBN ligand and VH032 as the VHL ligand. The synthetic routes and molecular characterization data for the final PROTAC compounds are provided in the Supplementary Information. This approach yielded multiple PROTAC molecules with linker lengths ranging from 1 to 4 polyethylene glycol (PEG) units, comprising compounds **20–22** (employing CRBN ligand) and **23–26** (employing VHL ligand).

**Scheme 1.**
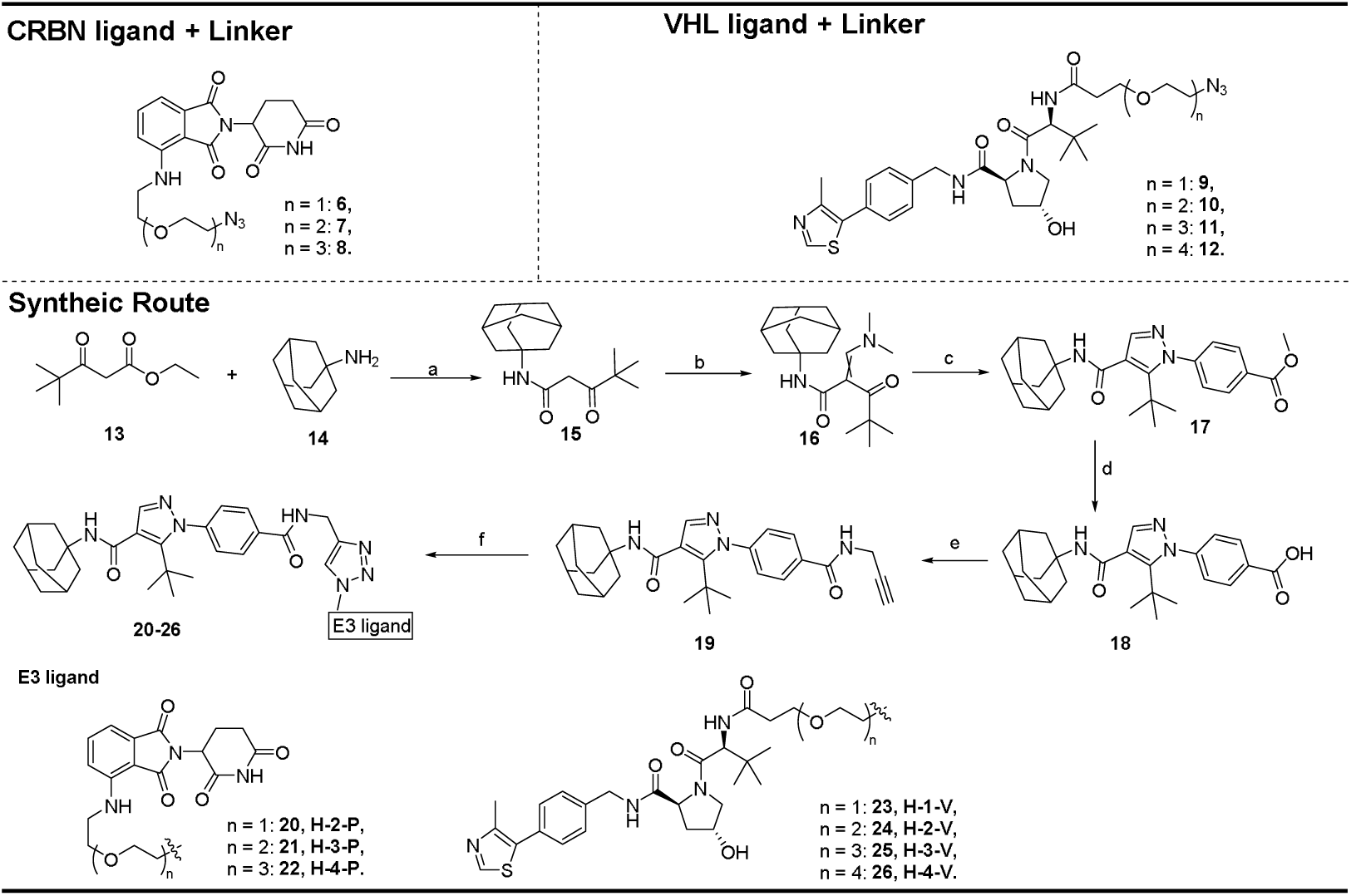
Synthetic route of PROTACs targeting 11β-HSD1 degradation. a) Xylene, 150 °C, 6 h; b) Dimethylformamide dimethyl acetal, 1,4-dioxane, reflux, 2 h; c) Methyl 4-hydrazineylbenzoate, acetic acid, ethanol, reflux, 1 h; d) LiOH, MeOH, HLO; e) HATU, TEA, DMF, DCM, 2 h; f) CuSOL, sodium ascorbate (VicNa), DMF.

**Scheme 2.**
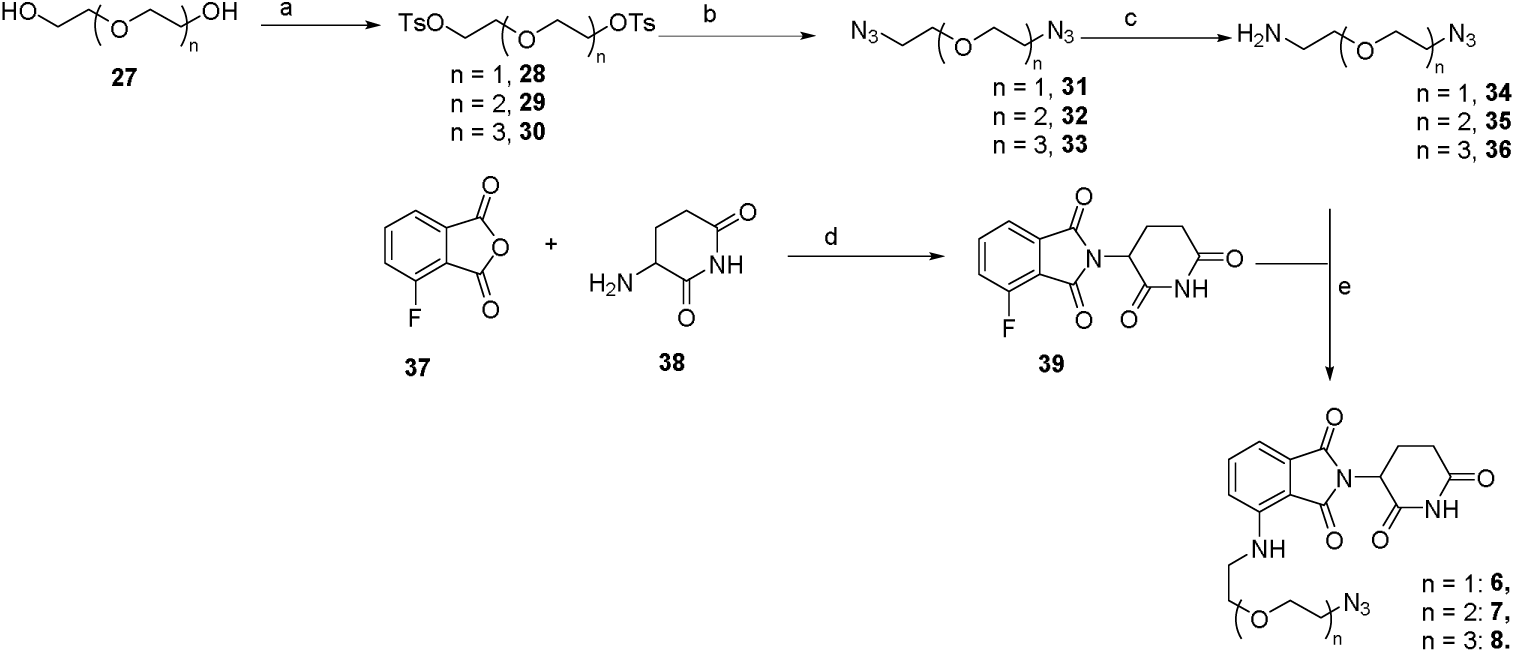
Synthetic route of CRBN ligand and linker. a) TsCl, Et_3_N, DCM, r.t. 8 h; b) TMSN_3_, TBAF, THF, 60°C, 6 h; c) PPh_3_, HCl, H_2_O/Ethyl acetate, r.t., 6 h; d) Et_3_N, THF, 80°C, 6 h; e) Et_3_N, DMF, 80L, 4 h

**Scheme 3.**
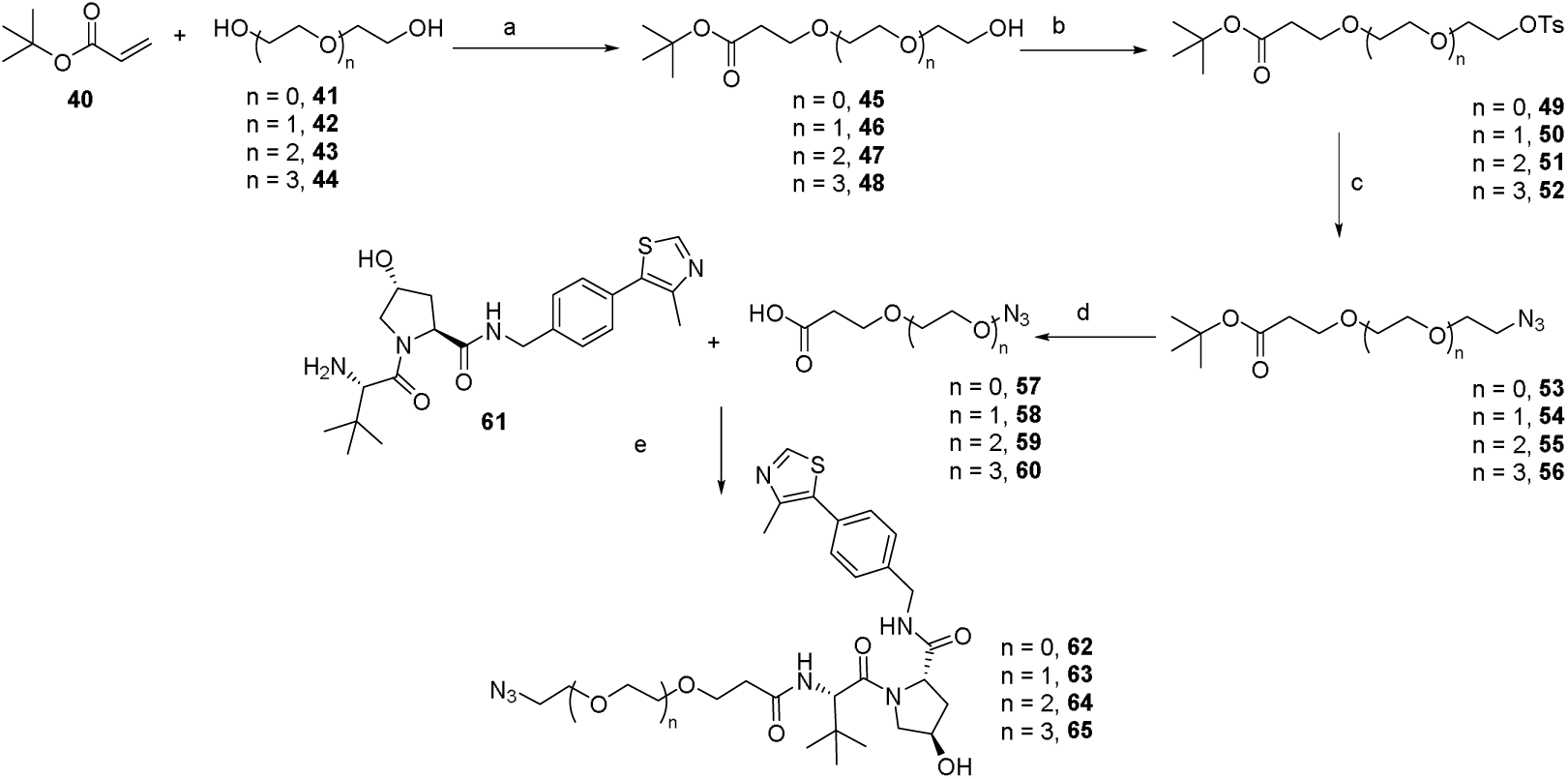
Synthetic route of VHL ligand and linker. a)Triton B, THF, 0°C->r.t., 6 h; b)TsCl, Et_3_N, DCM, 0°C->r.t., 6 h; c)TMSN_3_, TBAF, THF, 60°C, 6 h; d) HCl, DCM, r.t., 2 h; e) HATU, Et_3_N, DMF/DCM, r.t., 3 h.

**Figure 2.**
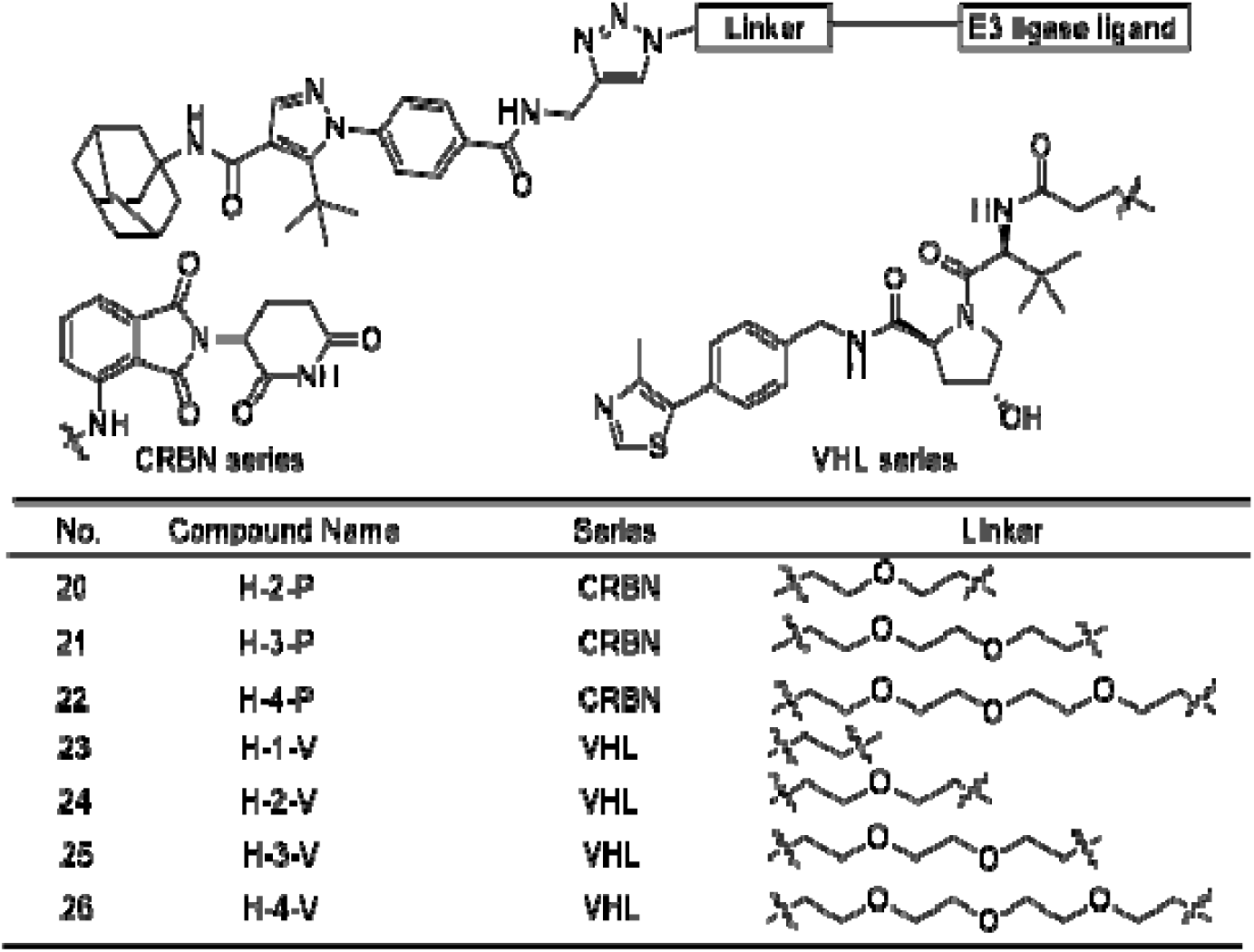
Chemical structure of degrader targeting 11β-HSD1

### 2.3 Degradation activity of synthesized PROTACs against 11β-HSD1

To evaluate the potency of degraders, we assessed the degradation activity of two series of molecules in cell lines. Following treatment, cells were harvested and lysed for subsequent western blot analysis. Notably, both CRBN- and VHL-based degraders induced concentration-dependent reductions in 11β-HSD1 levels, albeit with distinct potencies (**Figure 3A,B**). Degraders incorporating VHL ligands (**Figure 3B**) exhibited greater potency than those utilizing CRBN ligands (**Figure 3A**), despite the former PROTACs exhibiting significantly higher molecular weights. Compound H-3-V was selected for further evaluation as the most potent degrader (**Figure. 3C**). To quantify its degradation potency, cells were treated with increasing gradient concentrations of H-3-V, yielding a relative DC_50_ value of ∼1000 nM.

**Figure 3.**
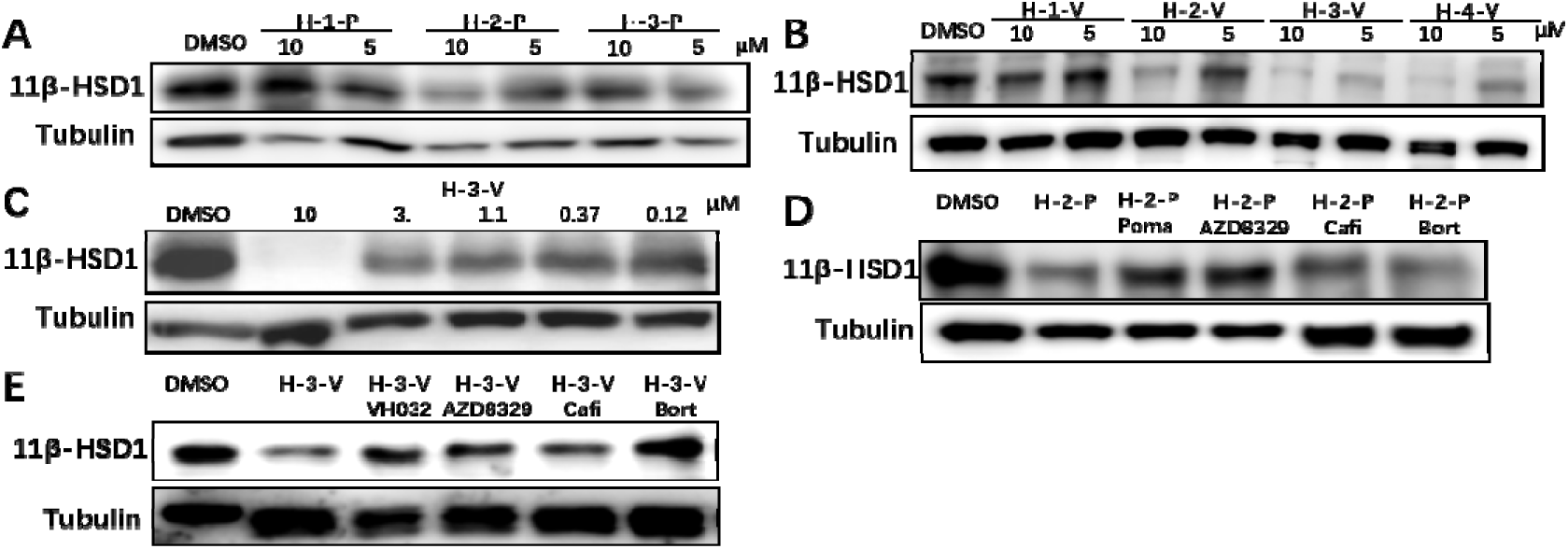
Evaluation of molecular activity by Western blotting. (A,B) Degradation of 11β-HSD1 by PROTAC molecule (C) Cellular-level assessment of H-3-V degradation activity demonstrating dose-dependent effects following treatment with varying concentrations of H-3-V. (A-C) Cells were treated for 48 hours. (D, E) Rescue assays. (D) Cells pretreated for 30 mins with 50 μM pomalidomide (Poma), 20 μM AZD8329, 80 nM carfilzomib (Cafi), or 200 nM bortezomib (Bort), subsequently treated with H-2-P for 12 hours, harvested, and analyzed by Western blotting. (E) Cells pretreated for 30 mins with 50 μM VH032, 20 μM AZD8329, 300 nM carfilzomib (Cafi), or 500 nM bortezomib (Bort), subsequently treated with H-3-V for 12 hours, harvested, and analyzed by Western blotting. (A-E) The MV-4-11 cell line was utilized throughout.

### 2.4 Study on degradation mechanism of H-2-P and H-3-V

We subsequently investigated whether this degradation resulted from PROTAC-induced protein ubiquitination. In subsequent experiments, H-2-P and H-3-V were selected as representative CRBN-(**Figure 3D**) and VHL-based (**Figure 3E**) compounds, respectively, for rescue experiments. Cells were pretreated with AZD8329, pomalidomide (for H-2-P), VH032 (for H-3-V), or carfilzomib. AZD8329, pomalidomide, and VH032 competitively bind to 11β-HSD1, CRBN, and VHL, respectively, thereby preventing ternary complex formation between 11β-HSD1, PROTAC, and the E3 ligase. Carfilzomib inhibits proteasomal activity. Consistent with expectations, pretreatment with each compound attenuated or prevented degradation, whereas H-2-P and H-3-V alone induced 11β-HSD1 degradation. These findings indicate that degradation is mediated by ternary complex formation and subsequent engagement of the ubiquitin-proteasome system (UPS), thereby confirming the fundamental mechanistic basis of PROTAC technology.

### 2.5 Study on the pharmacological effects in mouse models

Given the critical role of 11β-HSD1 in metabolic homeostasis and blood glucose regulation, we evaluated the *in vivo* efficacy of H-3-V in a high-fat diet (HFD)-induced type 2 diabetes mellitus (T2DM) mouse model. Intraperitoneal glucose tolerance test (IPGTT) revealed that two weeks of H-3-V treatment (i.p.) significantly improved glucose tolerance in diabetic mice (**Figure 4A,B**). These results suggest that H-3-V may enhance glucose-stimulated insulin secretion (GSIS), thereby contributing to its glucose-lowering effects. To further investigate whether H-3-V promotes GSIS, we performed an insulin secretion assay after two weeks of treatment. The result showed H-3-V administration significantly increased insulin secretion at 15 min post-glucose challenge, supporting its role in potentiating β-cell function (**Figure 4C, D**). Additionally, we monitored body weight and random blood glucose levels throughout the treatment period. H-3-V-treated mice exhibited a downward trend in both body weight and blood glucose compared to controls (**Figure 4E, F**). Our study demonstrates that H-3-V exerts significant pharmacological effects in HFD-induced T2DM mice, particularly in improving glucose tolerance and enhancing GSIS.

**Figure 4.**
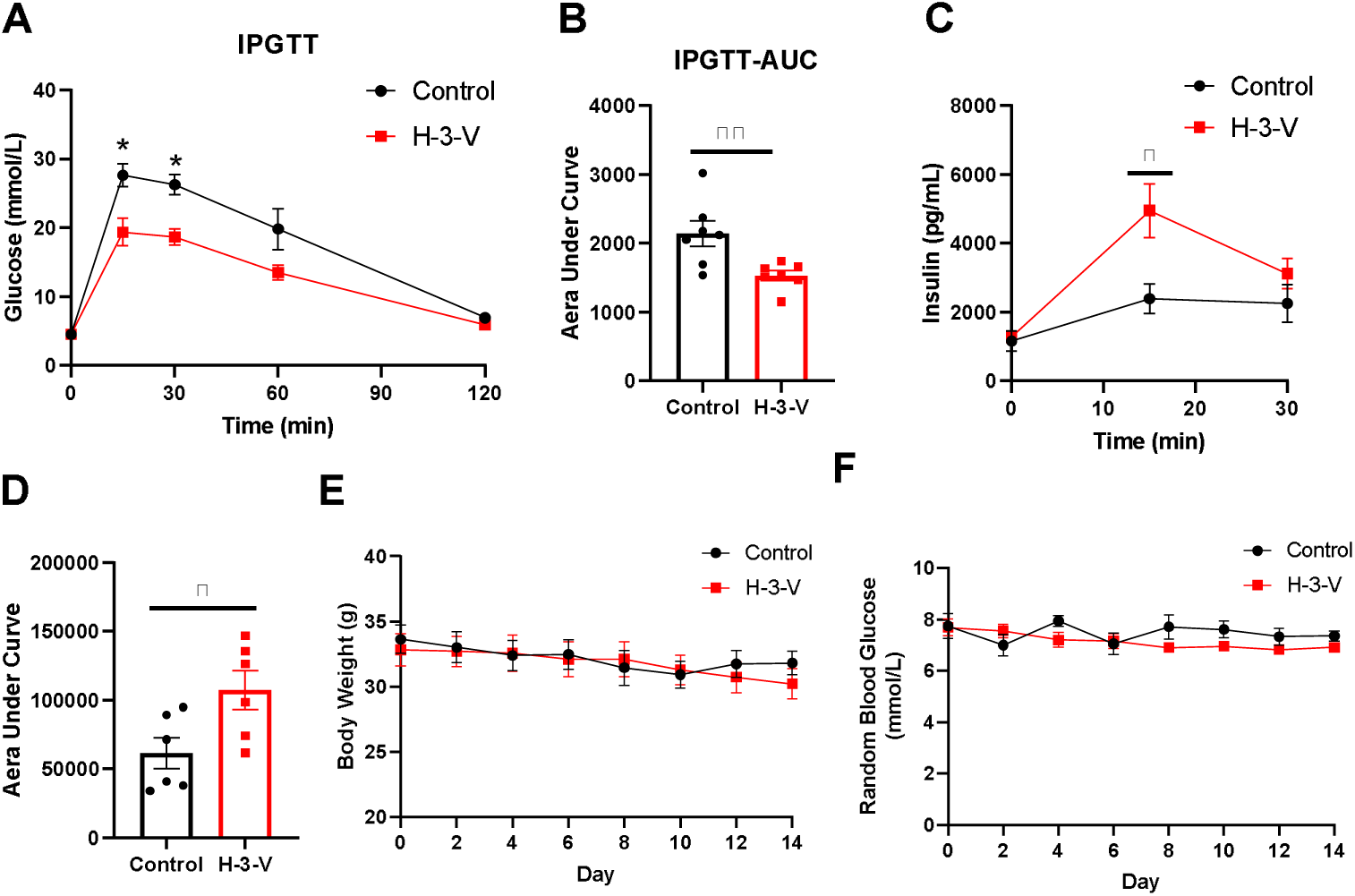
Administration of H-3-V decreased plasma insulin level, and improved glucose tolerance in high-fat fed mice. H-3-V (50 mg/kg) were administered intraperitoneal injection once every other day for 2 weeks to high fat diet-fed mice. (A) Effect of H-3-V on IPGTT (n=7). After the last treatment with H-3-V, mice were fasted overnight for 16 h and then subjected to intraperitoneal injection (i.p.) of D-glucose (2 g/kg). Blood glucose levels were measured at 0, 15, 30, 60, and 120 min post-injection. (B) The area under the curve (AUC) was calculated for (A). (C) Following the final administration of H-3-V, mice were fasted overnight for 16 h and received an i.p. injection of D-glucose (2 g/kg). Blood samples were collected from the inner canthus at 0, 15, and 30 min post-injection, and serum insulin levels were measured. (D) The AUC was calculated for (C). (E) Body weight of HFD mice with or without H-3-V for 2 weeks. (F) Random blood glucose HFD mice with or without H-3-V for 2 weeks. (\**P*L0.05, \*\**P*L0.01, compared with control group).

### 2.6 Molecular dynamics simulation of degrader-mediated molecular interactions

Although we have demonstrated through western blot that there is a significant difference in the activities of H-1-V and H-3-V, we still do not understand the source of this difference. Molecular dynamics stimulation (MD) can better illustrate the dynamic interaction process of the ternary complex. To elucidate the origin of this divergence, we conducted microsecond-scale MD simulations on systems containing H-1-V and H-3-V or without ligand (apo). The ternary complex structure comprising 11β-HSD1, the PROTAC molecule, and the VHL protein was initially constructed. After prolonged simulation, both the protein backbone and small-molecule geometries achieved essential stability, indicating convergence of the complex structures (**Supplementary Figure S2A**). Despite utilizing highly similar initial configurations, apo, H-1-V and H-3-V group produced markedly divergent simulation trajectories. Analysis of protein structural fluctuations revealed that the ternary complex with H-3-V exhibited greater conformational volatility than the H-1-V system (**Figure 5A**). Besides, the H-3-V complex mediated more substantial hydrogen-bond interactions between 11β-HSD1 and VHL (**Figure 5B**). Subsequently, we calculated the MM-GBSA of the trajectory frame by frame. The trajectory shows that each trajectory presents periodic changes and has reached a relatively stable state (**Supplementary Figure S2B**). Meanwhile, the presence of small molecules plays a significant role in the formation of ternary complexes. The systems without small molecules have significantly higher values on the ΔMM-GBSA index compared to those containing small molecules. Then, we computed the free energy landscapes (FELs) for both systems (**Supplementary Figure S2C**). Projecting the collective motions of the H-1-V and H-3-V ternary complexes and apo system onto identical principal components revealed distinct distributions along the first principal component. Each simulation system shows significant differences and exhibits clear clustering. We used the Gaussian Mixture Models (GMM) method to classify the trajectories (**Figure 5C**). Subsequently, we conducted ΔMM-GBSA energy analysis for each cluster. The results indicated that the binding energies of the H-3-V groups were superior to those of H-1-V, which was more conducive to the formation of the ternary complex (**Supplementary Figure S2D**). This might clarify why H-3-V is more active than H-1-V.

**Figure 5.**
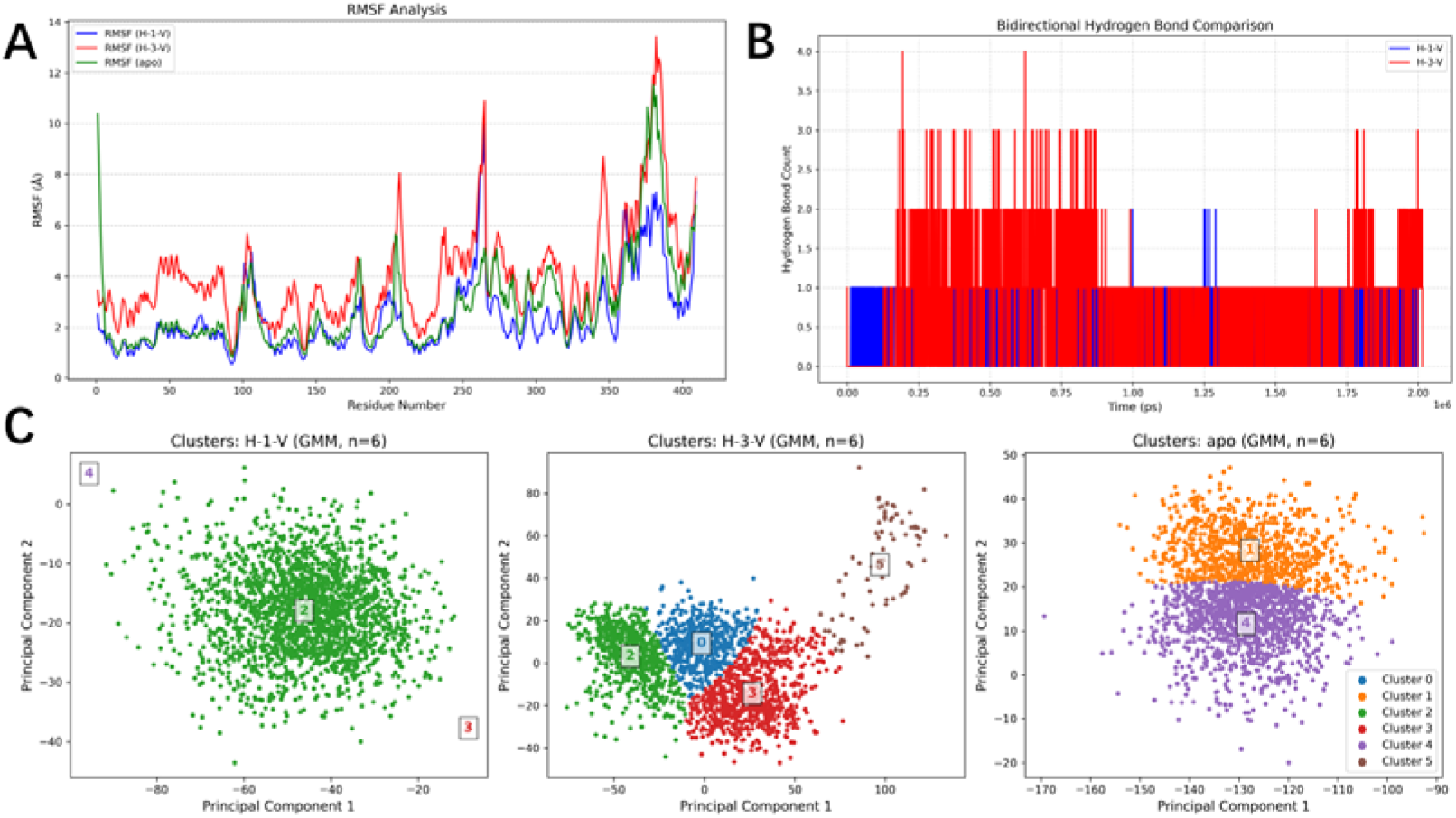
Molecular dynamics simulations were employed to assess the activity difference with H-1-V, H-3-V or without ligand (apo). (A) Root mean square fluctuation (RMSF) of protein atoms. (B) Number of hydrogen bond interactions between 11β-HSD1 and VHL. (C) Using the GMM classification method to cluster FEL.

## 3. Discussion

In this study, we report the development of a series of PROTACs targeting 11β-HSD1.Both CRBN-based and VHL-based PROTACs. effectively induce 11β-HSD1 degradation, albeit with varying potency. Furthermore, we elucidated the mechanism of action for compounds H-2-P and H-3-V by assessing their degradation efficacy under diverse conditions within the UPS. We subsequently conducted extensive *in vivo* verification of H-3-V activity, utilizing a diabetic metabolism model to confirm its pharmacological efficacy, thereby fully revealing the potential for this molecule’s *in vivo* application. H-3-V treatment significantly improved glucose tolerance and enhanced glucose-stimulated insulin secretion (GSIS) in HFD-induced T2DM mice, while showing only marginal effects on body weight and random blood glucose levels during the 2-week treatment period. These findings suggest that H-3-V may exert its antidiabetic effects primarily through improving β-cell function rather than through systemic metabolic alterations in the short term. MD simulation was used to explore the origin of the observed activity differences. Our results indicate that the ternary complex induced by H-3-V possesses a stronger binding free energy compared to that induced by H-1-V, alongside inducing a distinct binding mode. Concurrently, we acknowledge certain limitations inherent to this investigation. While longer linker chains are postulated to yield a greater number of conformations and potentially induce more favourable interactions, increased linker length inevitably incurs entropy loss during ternary complex formation, which may compromise degradation efficiency. Although we observed distinct free energy landscapes for H-3-V and H-1-V, this distinction was not fully exploited in the present work. In molecular dynamics simulations, we found that H-3-V exhibits some interaction phenomena similar to those of H-1-V clusters. However, H-3-V also generates some distinct cluster movements, and these differences in cluster movements might be the cause of the activity variations. It may be possible to combine cryo-electron microscopy to analyze the ternary complex model in the future, thereby providing assistance for rational design of PROTACs.

In summary, this work reports the development of a novel class of small molecule degraders targeting 11β-HSD1. These findings are anticipated to provide substantial support for elucidating the related functions and *in vivo* roles of 11β-HSD1.

## 4. Experimental Section

### 4.1. Chemistry

Reagents were procured from Macklin, Bidepharm, or Sigma-Aldrich. Solvents were obtained from Tansoole. Reactions were conducted using magnetic stirring in synthesis-grade glassware. Thin layer chromatography (TLC) was performed on silica gel-coated glass plates incorporating fluorescent indicators. Purification was achieved via column chromatography employing commercially available silica gel (300-400 mesh) supplied by Qingdao Haiyang Chemical Co., Ltd. (China). Nuclear magnetic resonance (NMR) spectra were acquired using a Bruker Avance 300 MHz spectrometer, utilizing tetramethylsilane (TMS) as the internal standard in DMSO-*d*6. High-resolution mass spectrometry (HR-MS) analyses were conducted on a waters SYNAPT G2 mass spectrometer.

#### 4.1.1. Synthetic route

The general procedure for synthesizing CRBN ligands 6-8 is detailed in the article published by Zixuan An et al[24].

Compound 40 (128 mg, 1.0 mmol) and the corresponding dihydroxy compounds (41–44, 5.0 mmol) were dissolved in dichloromethane (DCM, 10 mL) within a dry 25 mL round-bottom flask at ambient temperature. The reaction mixture was cooled to 0 °C under magnetic stirring. Triton B (0.05 mmol) was subsequently added. The reaction was then allowed to warm to room temperature. After stirring for 6 h, reaction completion was confirmed by LCMS analysis. The solvent was removed under reduced pressure. The crude product was purified by silica gel column chromatography to afford compounds 45–48 as colorless oils in yields ranging from 63% to 72%.

Compounds 45–48 (1.0 mmol) and tosyl chloride (210 mg, 1.1 mmol) were dissolved in dichloromethane (DCM, 10 mL) within a dry 25 mL round-bottom flask at 0 °C under magnetic stirring. EtLN (278 μL, 2.0 mmol) was subsequently added. After stirring for 1 h, the reaction was allowed to warm to room temperature. Stirring was continued for an additional 6 h, after which reaction completion was confirmed by LCMS analysis. The solvent was removed under reduced pressure. The crude product was purified by silica gel column chromatography to afford compounds 49–52 in yields of approximately 80%.

Compounds 49–52 (1.0 mmol) and azidotrimethylsilane (138 mg, 1.2 mmol) were dissolved in THF (10 mL) within a dry 25 mL round-bottom flask at room temperature. TBAF ((392 mg, 1.5 mmol) was subsequently added. The reaction mixture was then heated to 60 °C. After stirring for 6 h, reaction completion was confirmed by LCMS analysis. The solvent was removed under reduced pressure. The crude product was purified by silica gel column chromatography to afford compounds 53–56 as colorless oils in yields ranging from 55% to 69%.

Compounds 53–56 (1.0 mmol) were dissolved in DCM ((5 mL) within a dry 25 mL round-bottom flask at room temperature. 4 M HCl in dioxane (2 mL) was added. After stirring for 2 h, reaction completion was confirmed by LCMS analysis. The solvent was removed under reduced pressure. The crude material was employed directly in the subsequent step without further purification to yield compounds 57–60.

Compounds 57–60 (0.1 mmol), dissolved in N,N-dimethylformamide (DMF, 3 mL), were combined with compound 61 (430 mg, 0.1 mmol), N,N-diisopropylethylamine (DIPEA, 39 mg, 0.3 mmol), and O-(7-azabenzotriazol-1-yl)-N,N,N′,N′-tetramethyluronium hexafluorophosphate (HATU, 42 mg, 0.11 mmol) in a dry 10 mL single-necked flask at room temperature. The mixture was stirred at room temperature for 3 h. LCMS analysis confirmed reaction completion. The crude product was purified by silica gel column chromatography to afford compounds 62–65 as yellow oils in yields ranging from 65% to 69%.

##### Compound 15: N-(adamantan-1-yl)-4,4-dimethyl-3-oxopentanamide

Compound 13 (344 mg, 2 mmol) and compound 14 (302 mg, 2 mmol) were dissolved in xylene (10 mL) in a sealed tube. The reaction mixture was then placed under a nitrogen atmosphere and heated at 150 °C for 6 h. Thin-layer chromatography (TLC) indicated the complete consumption of starting materials. The crude product was purified by silica gel column chromatography to afford 487 mg of an off-white solid in 88% yield.

##### Compound 16: N-(adamantan-1-yl)-2-((dimethylamino)methylene)-4,4-dimethyl-3-oxopentanamide

Compound 15 (4.9 g, 17.68 mmol) and dimethylformamide acetal (2.32 g, 19.45 mmol) were dissolved in 20 mL of 1,4-dioxane within a 50 mL round-bottom flask at ambient temperature. Subsequently, the reaction mixture was refluxed for 6 h. Thin-layer chromatography (TLC) analysis confirmed the complete consumption of starting materials. The solvent was removed under reduced pressure, followed by washing the residue with 10 mL of a co-solvent mixture (EA/PE, 1:2 v/v). The resulting crude product (5.8 g) was collected by filtration as a precipitate without further purification.

##### Compound 17: methyl 4-(4-(adamantan-1-ylcarbamoyl)-5-(tert-butyl)-1H-pyrazol-1-yl)benzoate

Compound 16 (4 g, 12 mmol) and methyl 4-hydrazineylbenzoate hydrochloride (2.44 g, 12 mmol) were dissolved in ethanol (15 mL). Subsequently, acetic acid (10 drops) was added. The reaction mixture was heated at 80 °C for 2 h. Reaction completion was confirmed by thin-layer chromatography (TLC), indicating full consumption of the starting materials. The solvent was removed under reduced pressure. The crude product was purified by silica gel column chromatography, yielding 2.2 g of a yellow solid product (42% yield).

##### Compound 18: 4-(4-(adamantan-1-ylcarbamoyl)-5-(tert-butyl)-1H-pyrazol-1-yl)benzoic acid

Compound 17 (2 g, 4.6 mmol) was dissolved in 10 mL of co-solvent MeOH/H_2_O (3:1 v/v). LiOH (330 mg, 13.8 mmol) was added, and the resulting mixture was stirred overnight at ambient temperature. Reaction completion was confirmed by thin-layer chromatography (TLC), indicating complete consumption of the starting materials. The solvent was removed under reduced pressure. The crude product was purified by silica gel column chromatography, affording 1.73 g of a yellow solid product (89% yield).

##### Compound 19: N-(adamantan-1-yl)-5-(tert-butyl)-1-(4-(prop-2-yn-1-ylcarbamoyl)phenyl)-1H-pyrazole-4-ca rboxamide

Compounds 18 (421 mg, 1 mmol) dissolved in anhydrous N,N-dimethylformamide (DMF, 10 mL) were combined with prop-2-yn-1-amine ((66 mg, 1.2 mmol), N,N-diisopropylethylamine (DIPEA, 387 mg, 3 mmol), and O-(7-azabenzotriazol-1-yl)-N,N,N′,N′-tetramethyluronium hexafluorophosphate (HATU, 419 mg, 1.1 mmol) in a dry 25 mL single-necked flask at room temperature. The resulting mixture was stirred at this temperature for 3 hours. Reaction completion was confirmed by LCMS analysis. The crude product was purified by silica gel column chromatography, affording 391 mg of compound 19 as a white solid in 86% yield.

##### Compounds 20–26: General procedure for Click Reaction

Compound 19 (10 mg, 0.022 mmol, 1 equiv) and the corresponding E3 ligand (0.022 mmol, 1 equiv), synthesized as described in Sections 1.1.1 and 1.1.2, were dissolved in DMF (2 mL). A 1 M aqueous solution of CuSOL and a 1 M aqueous solution of sodium ascorbate (VicNa) were prepared. Subsequently, 0.05 equiv of the CuSOL solution and 0.10 equiv of the VicNa solution were added. The reaction mixture was stirred at ambient temperature for 12 h. Reaction completion was confirmed by LCMS analysis. The crude product was purified by silica gel column chromatography, affording the final compound as a white solid for the VH032-based derivative and as a yellow solid for the pomalidomide-based derivative in approximately 50% yield.

#### 4.1.2. NMR and and Mass Spectrometry Data for Compounds 20–26

##### N-(adamantan-1-yl)-5-(tert-butyl)-1-(4-(((1-(2-(2-((2-(2,6-dioxopiperidin-3-yl)-1,3-dioxoisoin dolin-4-yl)amino)ethoxy)ethyl)-1H-1,2,3-triazol-4-yl)methyl)carbamoyl)phenyl)-1H-pyrazole -4-carboxamide (20)

^1^H NMR (300 MHz, DMSO-*d*_6_) δ 11.10 (s, 1H), 9.27 – 9.12 (m, 1H), 8.07 – 7.91 (m, 3H), 7.72 (s, 1H), 7.63 – 7.50 (m, 2H), 7.44 (d, *J* = 8.1 Hz, 2H), 7.08 (dd, *J* = 19.6, 7.8 Hz, 2H), 6.60 (t, *J* = 5.8 Hz, 1H), 5.08 (dd, *J* = 12.5, 5.2 Hz, 1H), 4.63 – 4.44 (m, 4H), 3.86 (t, *J* = 5.3 Hz, 2H), 3.61 (d, *J* = 5.4 Hz, 2H), 3.52 – 3.38 (m, 2H), 2.96 – 2.82 (m, 1H), 2.65 – 2.54 (m, 2H), 2.11 – 1.95 (m, 11H), 1.65 (s, 5H), 1.19 (s, 9H).

^13^C NMR (75 MHz, DMSO) δ 172.78, 170.07, 168.92, 167.27, 165.23, 164.83, 148.99, 146.35, 144.69, 144.58, 138.83, 136.21, 134.72, 132.07, 128.01, 127.93, 123.46, 119.41, 117.41, 110.69, 109.25, 68.86, 68.70, 51.25, 49.23, 48.55, 41.54, 40.75, 36.11, 34.90, 33.03, 30.98, 30.45, 28.82, 22.14.

HRMS (m/z): calculated for C_45_H_53_N_10_O_7_ [M+H]^+^: 845.4099; found: 845.4108 calculated for C_45_H_52_N_10_O_7_Na [M+Na]^+^: 867.3918; found: 867.3950.

##### N-(adamantan-1-yl)-5-(tert-butyl)-1-(4-(((1-(2-(2-(2-((2-(2,6-dioxopiperidin-3-yl)-1,3-dioxoiso indolin-4-yl)amino)ethoxy)ethoxy)ethyl)-1H-1,2,3-triazol-4-yl)methyl)carbamoyl)phenyl)-1H -pyrazole-4-carboxamide (21)

^1^H NMR (300 MHz, DMSO-*d*_6_) δ 11.08 (s, 1H), 9.20 (t, *J* = 5.7 Hz, 1H), 8.03 – 7.93 (m, 3H), 7.70 (s, 1H), 7.63 – 7.49 (m, 2H), 7.48 – 7.39 (m, 2H), 7.12 (d, *J* = 8.6 Hz, 1H), 7.03 (d, *J* = 7.1 Hz, 1H), 6.64 – 6.54 (m, 1H), 5.05 (dd, *J* = 12.9, 5.2 Hz, 1H), 4.56 – 4.43 (m, 4H), 3.81 (t, *J* = 5.2 Hz, 2H), 3.61 – 3.49 (m, 6H), 3.44 (t, *J* = 5.3 Hz, 2H), 2.96 – 2.78 (m, 1H), 2.64 – 2.51 (m, 2H), 2.06 – 1.96 (m, 11H), 1.64 (s, 6H), 1.18 (s, 9H).

^13^C NMR (75 MHz, DMSO) δ 172.78, 170.07, 168.95, 167.27, 165.22, 164.83, 149.00, 146.38, 144.73, 144.60, 138.83, 136.22, 134.70, 132.07, 128.03, 127.92, 123.37, 119.41, 117.43, 110.68, 109.23, 69.62, 69.56, 68.84, 68.79, 51.26, 49.30, 48.54, 41.64, 40.76, 36.11, 34.92, 33.03, 30.97, 30.45, 28.83, 22.12.

HRMS (m/z): calculated for C_47_H_57_N_10_O_8_ [M+H]^+^: 889.4361; found: 889.4399 calculated for C_47_H_56_N_10_O_8_Na [M+Na]^+^: 911.4180; found: 911.4188.

##### N-(adamantan-1-yl)-5-(tert-butyl)-1-(4-(((1-(2-(2-(2-(2-((2-(2,6-dioxopiperidin-3-yl)-1,3-dioxo isoindolin-4-yl)amino)ethoxy)ethoxy)ethoxy)ethyl)-1H-1,2,3-triazol-4-yl)methyl)carbamoyl)phenyl)-1H-pyrazole-4-carboxamide (22)

^1^H NMR (300 MHz, DMSO) δ 11.09 (s, 1H), 9.20 (t, *J* = 5.8 Hz, 1H), 8.10 – 7.89 (m, 3H), 7.70 (s, 1H), 7.63 – 7.50 (m, 2H), 7.44 (d, *J* = 8.1 Hz, 2H), 7.13 (d, *J* = 8.6 Hz, 1H), 7.03 (d, *J* = 7.1 Hz, 1H), 6.59 (t, *J* = 5.8 Hz, 1H), 5.05 (dd, *J* = 12.8, 5.3 Hz, 1H), 4.57 – 4.43 (m, 4H), 3.79 (t, *J* = 5.3 Hz, 2H), 3.62 – 3.42 (m, 12H), 2.96 – 2.79 (m, 1H), 2.64 – 2.50 (m, 2H), 2.03 (s, 10H), 1.64 (s, 6H), 1.18 (s, 9H).

^13^C NMR (75 MHz, DMSO) δ 172.79, 170.06, 168.93, 167.28, 165.21, 164.83, 149.00, 146.39, 144.73, 144.61, 138.83, 136.21, 134.70, 132.08, 128.03, 127.93, 123.36, 119.42, 117.44, 110.66, 109.22, 69.74, 69.69, 69.53, 68.84, 68.72, 51.25, 49.27, 48.54, 41.67, 40.75, 36.10, 34.92, 33.04, 30.97, 30.45, 28.83, 22.13.

HRMS (m/z): calculated for C_49_H_61_N_10_O_9_ [M+H]^+^: 933.4623; found: 933.4659 calculated for C_49_H_60_N_10_O_9_Na [M+Na]^+^: 955.4442; found: 955.4392.

##### N-(adamantan-1-yl)-5-(tert-butyl)-1-(4-(((1-(2-(3-(((S)-1-((2S,4R)-4-hydroxy-2-((4-(4-methylthiazol-5-yl)benzyl)carbamoyl)pyrrolidin-1-yl)-3,3-dimethyl-1-oxobutan-2-yl)amino)-3-oxopropoxy)ethyl)-1H-1,2,3-triazol-4-yl)methyl)carbamoyl)phenyl)-1H-pyrazole-4-carboxamide (23)

^1^H NMR (300 MHz, DMSO) δ 9.19 (t, *J* = 5.7 Hz, 1H), 8.98 (s, 1H), 8.57 (t, *J* = 6.0 Hz, 1H), 8.04 – 7.90 (m, 4H), 7.71 (s, 1H), 7.52 (s, 1H), 7.49 – 7.32 (m, 6H), 5.14 (t, *J* = 3.6 Hz, 1H), 4.58 – 4.17 (m, 9H), 3.84 – 3.53 (m, 6H), 2.55 (d, *J* = 6.5 Hz, 1H), 2.44 (d, *J* = 1.4 Hz, 3H), 2.36 (dt, *J* = 13.9, 6.2 Hz, 1H), 2.03 (s, 10H), 1.96 – 1.83 (m, 1H), 1.64 (s, 6H), 1.19 (s, 9H), 0.92 (s, 9H).

^13^C NMR (75 MHz, DMSO) δ 171.87, 169.78, 169.53, 165.23, 164.81, 151.39, 148.99, 147.69, 144.65, 144.59, 139.45, 138.81, 134.71, 131.13, 129.62, 128.60, 128.00, 127.93, 127.39, 123.37, 119.40, 68.84, 68.40, 66.74, 58.68, 56.35, 51.25, 49.23, 41.62, 40.75, 37.93, 36.09, 35.42, 35.27, 35.13, 34.91, 33.02, 30.45, 28.81, 26.31, 15.91.

HRMS (m/z): calculated for C_55_H_72_N_11_O_7_S [M+H]^+^: 1030.5337; found: 1030.5295 calculated forC_55_H_71_N_11_O_7_SNa [M+Na]^+^: 1052.5156; found: 1052.5198.

##### N-(adamantan-1-yl)-5-(tert-butyl)-1-(4-(((1-(2-(2-(3-(((S)-1-((2S,4R)-4-hydroxy-2-((4-(4-methylthiazol-5-yl)benzyl)carbamoyl)pyrrolidin-1-yl)-3,3-dimethyl-1-oxobutan-2-yl)amino)-3-oxo propoxy)ethoxy)ethyl)-1H-1,2,3-triazol-4-yl)methyl)carbamoyl)phenyl)-1H-pyrazole-4-carboxamide (24)

^1^H NMR (300 MHz, DMSO) δ 9.21 (s, 1H), 8.98 (s, 1H), 8.56 (s, 1H), 8.05 – 7.87 (m, 4H), 7.71 (s, 1H), 7.53 (s, 1H), 7.50 – 7.35 (m, 6H), 5.19 – 5.08 (m, 1H), 4.59 – 4.32 (m, 8H), 4.29 – 4.17 (m, 1H), 3.79 (t, *J* = 5.2 Hz, 2H), 3.68 – 3.40 (m, 8H), 2.56 (s, 1H), 2.44 (s, 3H), 2.41 – 2.27 (m, 1H), 2.03 (s, 10H), 1.90 (s, 1H), 1.64 (s, 6H), 1.19 (s, 9H), 0.93 (s, 9H).

^13^C NMR (75 MHz, DMSO) δ 171.90, 169.94, 169.52, 165.25, 164.84, 151.44, 149.00, 147.71, 144.72, 144.60, 139.48, 138.82, 134.72, 131.14, 129.63, 128.63, 128.03, 127.95, 127.41, 123.38, 119.41, 69.43, 69.36, 68.86, 68.75, 66.90, 58.70, 56.36, 56.29, 51.25, 49.26, 41.64, 40.75, 37.96, 36.10, 35.62, 35.35, 34.93, 33.04, 30.46, 28.82, 26.31, 15.93.

HRMS (m/z): calculated for C_57_H_76_N_11_O_8_S [M+H]^+^: 1074.5599; found: 1074.5560 calculated forC_57_H_75_N_11_O_8_SNa [M+Na]^+^: 1096.5418; found: 1096.5460.

##### N-(adamantan-1-yl)-5-(tert-butyl)-1-(4-(((1-((S)-14-((2S,4R)-4-hydroxy-2-((4-(4-methylthiazol-5-yl)benzyl)carbamoyl)pyrrolidine-1-carbonyl)-15,15-dimethyl-12-oxo-3,6,9-trioxa-13-azah exadecyl)-1H-1,2,3-triazol-4-yl)methyl)carbamoyl)phenyl)-1H-pyrazole-4-carboxamide (25)

^1^H NMR (300 MHz, DMSO) δ 9.26 – 9.15 (m, 1H), 8.98 (s, 1H), 8.56 (t, *J* = 6.0 Hz, 1H), 8.04 – 7.88 (m, 4H), 7.71 (s, 1H), 7.53 (s, 1H), 7.49 – 7.34 (m, 6H), 5.13 (d, *J* = 3.4 Hz, 1H), 4.60 – 4.31 (m, 8H), 4.22 (dd, *J* = 16.0, 5.4 Hz, 1H), 3.80 (t, *J* = 5.3 Hz, 2H), 3.71 – 3.40 (m, 12H), 2.59 – 2.52 (m, 1H), 2.41 – 2.28 (m, 1H), 2.01 (s, 10H), 1.96 – 1.83 (m, 1H), 1.64 (s, 6H), 1.19 (s, 9H), 0.93 (s, 9H).

^13^C NMR (75 MHz, DMSO) δ 171.90, 169.92, 169.52, 165.21, 164.84, 151.44, 149.00, 147.70, 144.73, 144.60, 139.49, 138.83, 134.71, 131.15, 129.63, 128.63, 128.03, 127.93, 127.41, 123.37, 119.41, 69.67, 69.63, 69.49, 69.45, 68.85, 68.73, 66.92, 58.69, 56.35, 56.28, 51.26, 49.27, 41.63, 40.75, 37.93, 36.10, 35.63, 35.35, 34.93, 33.04, 30.46, 28.82, 26.32, 15.93.

HRMS (m/z): calculated for C_59_H_80_N_11_O_9_S [M+H]^+^: 1118.5861; found: 1118.5835 calculated forC_59_H_79_N_11_O_9_SNa [M+Na]^+^: 1140.5681; found: 1140.5750.

##### N-(adamantan-1-yl)-5-(tert-butyl)-1-(4-(((1-((S)-17-((2S,4R)-4-hydroxy-2-((4-(4-methylthiazol-5-yl)benzyl)carbamoyl)pyrrolidine-1-carbonyl)-18,18-dimethyl-15-oxo-3,6,9,12-tetraoxa-16 -azanonadecyl)-1H-1,2,3-triazol-4-yl)methyl)carbamoyl)phenyl)-1H-pyrazole-4-carboxamide (26)

^1^H NMR (300 MHz, DMSO) δ 9.21 (s, 1H), 8.98 (s, 1H), 8.55 (s, 1H), 8.02 – 7.88 (m, 4H), 7.71 (s, 1H), 7.53 (s, 1H), 7.49 – 7.34 (m, 6H), 5.13 (d, *J* = 3.3 Hz, 1H), 4.60 – 4.31 (m, 8H), 4.22 (dd, *J* = 15.5, 5.2 Hz, 1H), 3.80 (t, *J* = 5.3 Hz, 2H), 3.72 – 3.37 (m, 16H), 2.60 – 2.52 (m, 1H), 2.44 (s, 3H), 2.40 – 2.26 (m, 1H), 2.03 (s, 10H), 1.96 – 1.82 (m, 1H), 1.64 (s, 6H), 1.19 (s, 9H), 0.93 (s, 9H).

^13^C NMR (75 MHz, DMSO) δ 171.91, 169.91, 169.52, 165.21, 164.84, 151.44, 148.99, 147.71, 144.72, 144.61, 139.49, 138.83, 134.71, 131.14, 129.63, 128.63, 128.03, 127.93, 127.41, 123.37, 119.41, 69.73, 69.67, 69.62, 69.51, 69.44, 68.86, 68.74, 66.93, 58.70, 56.36, 56.27, 51.26, 49.27, 41.64, 40.75, 37.93, 36.11, 35.64, 35.35, 34.93, 33.04, 30.46, 28.82, 26.32, 15.93.

HRMS (m/z): calculated for C_61_H_84_N_11_O_10_S [M+H]^+^: 1162.6123; found: 1162.6167 calculated forC_61_H_83_N_11_O_10_SNa [M+Na]^+^: 1184.5943; found: 1184.5989.

### 4.2. Cell line and culture

The human biphenotypic B myelomonocytic leukemia cell line MV4-11 was obtained from Procell system (CL-0572). Cells were cultured in Iscove’s Modified Dulbecco’s Medium (IMDM) (Gibco, C12440500BT) supplemented with 10% fetal bovine serum (FBS) (CELLiGENT, China, Cat#:CG1126B) and 1% penicillin-streptomycin (P/S) (AbbKine, USA, Cat#: BMC1030). Cells were maintained in sterile culture flasks at 37°C in a humidified incubator with 5% COL. The medium was refreshed every 48 hours to maintain optimal nutrient conditions. Cell density was routinely monitored and maintained between 1×10L and 1×10L cells/mL by subculturing or dilution with fresh medium. For cell treatment, MV4-11 cells were seeded in 6-well plates at a density of 5 x 10^6^ cells per well. After 24-hour incubation, cells were treated with the following compounds, drug or vehicle controls such as H-1-P, H-2-P, H-3-P, H-1-V, H-2-V, H-3-V, H-4-V, pomalidomide (bidepharm, BD235626), VH032 (bidepharm, BD00758812), bortezomib (bidepharm, BD106087), carfilzomib (bidepharm, BD292920), compound 18 and DMSO, for specified durations. After treatment, cells were lysed and subjected to Western blot analysis to determine 11β-HSD1 protein expression.

### 4.3. Animal

6 week old male C57BL/6J wild-type mice were obtained from Beijing Vital River Laboratory Animal Technology Co., Ltd. (Beijing, China). Mice were housed under a 12:12-hour light-dark cycle with free access to water. Prior to the study, all animals were acclimatized in the laboratory for at least two weeks. All animal care and experimental procedures were approved by the Animal Policy and Welfare Committee of Capital Medical University and conducted in accordance with the approved protocol.

After 8 weeks of high-fat diet (HFD, 60% fat, 20% carbohydrate, and 20% protein in calorie percentage; Research Diets) feeding, the mice were randomly divided into two groups (n=7 per group): (1) the HFD + vehicle control group and (2) the HFD + H-3-V group (a PROTAC degrader targeting 11β-HSD1, 50 mg/kg via intraperitoneal injection every other day). The treatment period lasted for two weeks, during which body weight and random blood glucose levels were monitored every other day.

### 4.4. Intraperitoneal glucose tolerance test (IPGTT) and intraperitoneal insulin resistance test (IPIRT) experiment

After a 16-hour overnight fast, mice underwent an intraperitoneal glucose tolerance test (IPGTT) and intraperitoneal insulin resistance test (IPIRT). For the IPGTT, mice received an intraperitoneal (i.p.) injection of glucose (2 g/kg body weight). Blood glucose levels were measured from tail vein blood at 0, 15, 50, 60, and 120 min post-injection using an ACCU-CHEK Performa glucometer (Roche Diagnostics, Basel, Switzerland).

For the IPIRT, blood samples were collected via the retro-orbital sinus at 0, 15, and 30 min after glucose administration. Serum was isolated by centrifugation, and insulin concentrations were quantified using a commercially available ELISA kit (NOVUS BIOLOGICALS, Littleton, CO, USA) according to the manufacturer’s protocol.

### 4.5. Western Blotting Analysis

Cells were lysed in freshly prepared RIPA buffer (Beyotime, 50 mM Tris-HCl pH 7.4, 150 mM NaCl, 1% NP-40, 0.5% sodium deoxycholate, 0.1% SDS) supplemented with 1 mM PMSF and protease inhibitors on ice for 30 min, followed by centrifugation at 12,000 × g for 10 min at 4°C. Protein concentrations were determined by BCA assay, and equal amounts (20–50 μg) were separated by SDS-PAGE and transferred to PVDF membranes. After blocking with 5% non-fat milk in TBST for 30 min, membranes were incubated overnight at 4°C with primary antibodies against 11β-HSD1 (Rabbit, abcam, ab157223, 1:1000), β-actin (Rabbit, Proteintech, 1:2000), or α-tubulin (Mouse, Proteintech, 1:2000), followed by HRP-conjugated secondary antibodies (1:1000, Beyotime, A0216, A0208) and ECL detection. Protein expression was quantified using ImageJ, with β-actin or α-tubulin as loading controls.

### 4.6. Statistical Analysis

All statistical analyses were performed using GraphPad Prism software (version 8.0; GraphPad Software, San Diego, CA). Quantitative data are presented as mean ± standard error of the mean (SEM) from at least three independent experiments. Between-group comparisons were analyzed using unpaired two-tailed Student’s t-tests. A probability value of *P* < 0.05 was considered statistically significant.

## Acknowledgment

This project was supported by the Beijing Municipal Natural Science Foundation (7244475). Beijing Hospitals Authority Innovation Studio of Young Staff Funding SupportLcode: 202105. We express our gratitude to Qianlong Liu from Tsinghua University for providing certain assistance in the establishment of the docking model. The author(s) used Kingsoft WPS AI tools in order to polish the language and improve readability.

## CRediT authorship contribution statement

**Liguo Wang:** Writing original draft, Project administration, Investigation, Conceptualization, supervision, Funding acquisition; **Juanjuan Zhu:** Writing original draft, Validation, supervision; **Xue Tao:** Validation, Formal analysis; **Yu Lu, Yuji Wang**: Formal analysis; **Ming He:** Software, Formal analysis.

## Data availability

All data supporting the findings of this study are available in the manuscript or supplementary material. Supplementary material to this article can be found online at XXXXXX.

## Reference

[1] H. Ma, G.Y. Sui, J.S. Park, F. Wang, Y. Ma, D.S. Shin, N. Rustamov, J.S. Jang, S.I. Chang, J. Lee, Y.S. Roh, Blockade of 11beta-hydroxysteroid dehydrogenase type 1 ameliorates metabolic dysfunction-associated steatotic liver disease and fibrosis, Heliyon 10(20) (2024) e39534.

[2] K. Chapman, M. Holmes, J. Seckl, 11beta-hydroxysteroid dehydrogenases: intracellular gate-keepers of tissue glucocorticoid action, Physiol. Rev. 93(3) (2013) 1139–206.

[3] T.C. Sandeep, J.L. Yau, A.M. MacLullich, J. Noble, I.J. Deary, B.R. Walker, J.R. Seckl, 11Beta-hydroxysteroid dehydrogenase inhibition improves cognitive function in healthy elderly men and type 2 diabetics, Proc. Natl. Acad. Sci. U. S. A. 101(17) (2004) 6734–9.

[4] N. Draper, P.M. Stewart, 11beta-hydroxysteroid dehydrogenase and the pre-receptor regulation of corticosteroid hormone action, J. Endocrinol. 186(2) (2005) 251–71.

[5] S.L. Lightman, M.T. Birnie, B.L. Conway-Campbell, Dynamics of ACTH and Cortisol Secretion and Implications for Disease, Endocr. Rev. 41(3) (2020).

[6] S.A. Morgan, E.L. McCabe, L.L. Gathercole, Z.K. Hassan-Smith, D.P. Larner, I.J. Bujalska, P.M. Stewart, J.W. Tomlinson, G.G. Lavery, 11beta-HSD1 is the major regulator of the tissue-specific effects of circulating glucocorticoid excess, Proc. Natl. Acad. Sci. U. S. A. 111(24) (2014) E2482–91.

[7] X. Li, J. Wang, Q. Yang, S. Shao, 11beta-Hydroxysteroid Dehydrogenase Type 1 in Obese Subjects With Type 2 Diabetes Mellitus, Am. J. Med. Sci. 354(4) (2017) 408–414.

[8] H. Masuzaki, H. Yamamoto, C.J. Kenyon, J.K. Elmquist, N.M. Morton, J.M. Paterson, H. Shinyama, M.G. Sharp, S. Fleming, J.J. Mullins, J.R. Seckl, J.S. Flier, Transgenic amplification of glucocorticoid action in adipose tissue causes high blood pressure in mice, J. Clin. Invest. 112(1) (2003) 83–90.

[9] T.M. Anil, A. Dandu, K. Harsha, J. Singh, N. Shree, V.S. Kumar, M.N. Lakshmi, V. Sunil, C. Harish, G.V. Balamurali, B.S. Naveen Kumar, A.S. Gopala, S. Pratibha, M. Sadasivuni, M.O. Anup, Y. Moolemath, M.V. Venkataranganna, M.R. Jagannath, B.P. Somesh, A novel 11beta-hydroxysteroid dehydrogenase type1 inhibitor CNX-010-49 improves hyperglycemia, lipid profile and reduces body weight in diet induced obese C57B6/J mice with a potential to provide cardio protective benefits, BMC Pharmacol Toxicol 15 (2014) 43.

[10] Y. Kotelevtsev, M.C. Holmes, A. Burchell, P.M. Houston, D. Schmoll, P. Jamieson, R. Best, R. Brown, C.R.W. Edwards, J.R. Seckl, J.J. Mullins, 11β-Hydroxysteroid dehydrogenase type 1 knockout mice show attenuated glucocorticoid-inducible responses and resist hyperglycemia on obesity orLJstress, Proceedings of the National Academy of Sciences 94(26) (1997) 14924–14929.

[11] S.D. Reichardt, A. Amouret, C. Muzzi, S. Vettorazzi, J.P. Tuckermann, F. Luhder, H.M. Reichardt, The Role of Glucocorticoids in Inflammatory Diseases, Cells 10(11) (2021).

[12] D. Kupczyk, R. Studzinska, R. Kolodziejska, S. Baumgart, M. Modrzejewska, A. Wozniak, 11beta-Hydroxysteroid Dehydrogenase Type 1 as a Potential Treatment Target in Cardiovascular Diseases, J Clin Med 11(20) (2022).

[13] L. Wang, J. Liu, A. Zhang, P. Cheng, X. Zhang, S. Lv, L. Wu, J. Yu, W. Di, J. Zha, X. Kong, H. Qi, Y. Zhong, G. Ding, BVT.2733, a selective 11beta-hydroxysteroid dehydrogenase type 1 inhibitor, attenuates obesity and inflammation in diet-induced obese mice, PLoS One 7(7) (2012) e40056.

[14] S. Shah, A. Hermanowski-Vosatka, K. Gibson, R.A. Ruck, G. Jia, J. Zhang, P.M. Hwang, N.W. Ryan, R.B. Langdon, P.U. Feig, Efficacy and safety of the selective 11beta-HSD-1 inhibitors MK-0736 and MK-0916 in overweight and obese patients with hypertension, J. Am. Soc. Hypertens. 5(3) (2011) 166–76.

[15] J. Rosenstock, S. Banarer, V.A. Fonseca, S.E. Inzucchi, W. Sun, W. Yao, G. Hollis, R. Flores, R. Levy, W.V. Williams, J.R. Seckl, R. Huber, I.P. Investigators, The 11-beta-hydroxysteroid dehydrogenase type 1 inhibitor INCB13739 improves hyperglycemia in patients with type 2 diabetes inadequately controlled by metformin monotherapy, Diabetes Care 33(7) (2010) 1516–22.

[16] X.Y. Ye, S.Y. Chen, S. Wu, D.S. Yoon, H. Wang, Z. Hong, S.P. O’Connor, J. Li, J.J. Li, L.J. Kennedy, S.J. Walker, A. Nayeem, S. Sheriff, D.M. Camac, V. Ramamurthy, P.E. Morin, R. Zebo, J.R. Taylor, N.N. Morgan, R.P. Ponticiello, T. Harrity, A. Apedo, R. Golla, R. Seethala, M. Wang, T.W. Harper, B.G. Sleczka, B. He, M. Kirby, D.K. Leahy, J. Li, R.L. Hanson, Z. Guo, Y.X. Li, J.D. DiMarco, R. Scaringe, B. Maxwell, F. Moulin, J.C. Barrish, D.A. Gordon, J.A. Robl, Discovery of Clinical Candidate 2-((2S,6S)-2-Phenyl-6-hydroxyadamantan-2-yl)-1-(3’-hydroxyazetidin-1-yl)ethanone [BMS-816336], an Orally Active Novel Selective 11beta-Hydroxysteroid Dehydrogenase Type 1 Inhibitor, J. Med. Chem. 60(12) (2017) 4932–4948.

[17] P. Morentin Gutierrez, A. Gyte, J. deSchoolmeester, P. Ceuppens, J. Swales, C. Stacey, J.W. Eriksson, M. Sjostrand, C. Nilsson, B. Leighton, Continuous inhibition of 11beta-hydroxysteroid dehydrogenase type I in adipose tissue leads to tachyphylaxis in humans and rats but not in mice, Br J Pharmacol 172(20) (2015) 4806–16.

[18] A.G. Atanasov, L.G. Nashev, L. Gelman, B. Legeza, R. Sack, R. Portmann, A. Odermatt, Direct protein-protein interaction of 11beta-hydroxysteroid dehydrogenase type 1 and hexose-6-phosphate dehydrogenase in the endoplasmic reticulum lumen, Biochim Biophys Acta 1783(8) (2008) 1536–43.

[19] F. Yao, L. Chen, Z. Fan, F. Teng, Y. Zhao, F. Guan, M. Zhang, Y. Liu, Interplay between H6PDH and 11beta-HSD1 implicated in the pathogenesis of type 2 diabetes mellitus, Bioorg Med Chem Lett 27(17) (2017) 4107–4113.

[20] C. Cao, M. He, L. Wang, Y. He, Y. Rao, Chemistries of bifunctional PROTAC degraders, Chem. Soc. Rev. 51(16) (2022) 7066–7114.

[21] J.S. Scott, J. deSchoolmeester, E. Kilgour, R.M. Mayers, M.J. Packer, D. Hargreaves, S. Gerhardt, D.J. Ogg, A. Rees, N. Selmi, A. Stocker, J.G. Swales, P.R. Whittamore, Novel acidic 11beta-hydroxysteroid dehydrogenase type 1 (11beta-HSD1) inhibitor with reduced acyl glucuronide liability: the discovery of 4-[4-(2-adamantylcarbamoyl)-5-tert-butyl-pyrazol-1-yl]benzoic acid (AZD8329), J. Med. Chem. 55(22) (2012) 10136–47.

[22] T. Cheng, Y. Zhao, X. Li, F. Lin, Y. Xu, X. Zhang, Y. Li, R. Wang, L. Lai, Computation of octanol-water partition coefficients by guiding an additive model with knowledge, J. Chem. Inf. Model. 47(6) (2007) 2140–8.

[23] A.R. Alanzi, A.Y. Moussa, M.S. Alsalhi, T. Nawaz, I. Ali, Integration of pharmacophore-based virtual screening, molecular docking, ADMET analysis, and MD simulation for targeting EGFR: A comprehensive drug discovery study using commercial databases, PLoS One 19(12) (2024) e0311527.

[24] Z. An, W. Lv, S. Su, W. Wu, Y. Rao, Developing potent PROTACs tools for selective degradation of HDAC6 protein, Protein Cell 10(8) (2019) 606–609.

