## Supplementary material for "Design, synthesis, and pharmacological evaluation of novel PROTAC degraders targeting 11β-HSD1 for metabolic disease intervention": Supplementary Information-submit.docx

**Supplementary Figures**

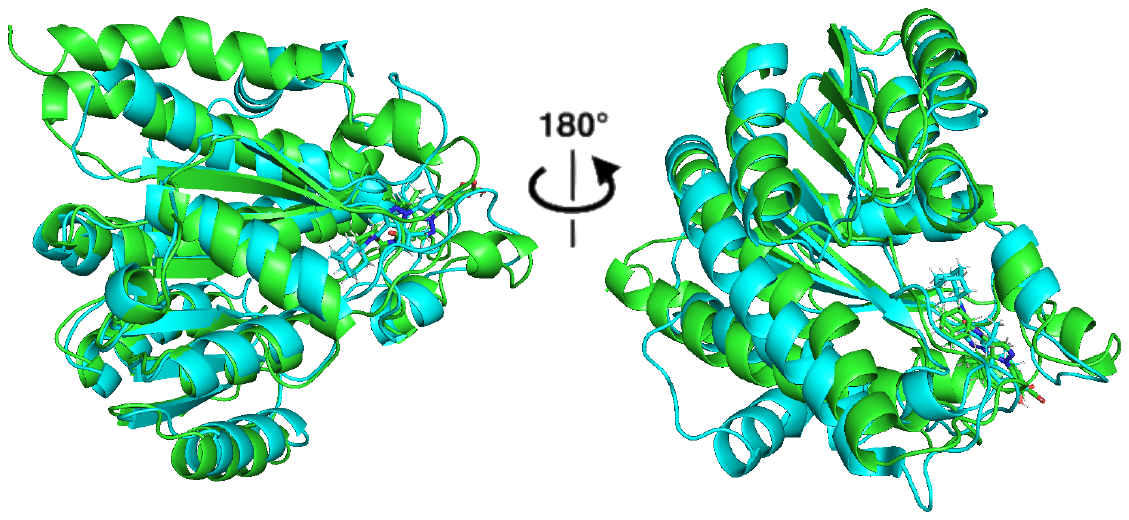

**Figure S1.** Stable molecular dynamics binding model. AZD8329 and the modified molecule 1-adamantyl-substituted molecule were docked into 11β-HSD1, and molecular dynamics simulations were conducted. After stabilization, the last frame was taken for protein alignment.

**Figure S2.** Molecular dynamics simulations were employed to assess the activity difference with H-1-V, H-3-V or without ligand (apo). (A) Root mean square deviation (RMSD) of the protein backbone and non-hydrogen atoms. (B) Calculate the MMGBSA value of the trajectory frame by frame. (C) Free energy landscape (FEL) generated by projecting the H-1-V and apo simulation data onto the principal component space defined by H-3-V. (D) The MM-GBSA method is used to conduct free energy statistical analysis on each cluster.

**Material and methods**

**1 Chemistry**

Reagents were procured from Macklin, Bidepharm, or Sigma-Aldrich. Solvents were obtained from Tansoole. Reactions were conducted using magnetic stirring in synthesis-grade glassware. Thin layer chromatography (TLC) was performed on silica gel-coated glass plates incorporating fluorescent indicators. Purification was achieved via column chromatography employing commercially available silica gel (300-400 mesh) supplied by Qingdao Haiyang Chemical Co., Ltd. (China). Nuclear magnetic resonance (NMR) spectra were acquired using a Bruker Avance 300 MHz spectrometer, utilizing tetramethylsilane (TMS) as the internal standard in DMSO-*d*6. High-resolution mass spectrometry (HR-MS) analyses were conducted on a waters SYNAPT G2 mass spectrometer.

**1.1 Synthetic route**

1.1.1 The general procedure for synthesizing CRBN ligands 6-8 is detailed in the article published by Zixuan An et al^1^.

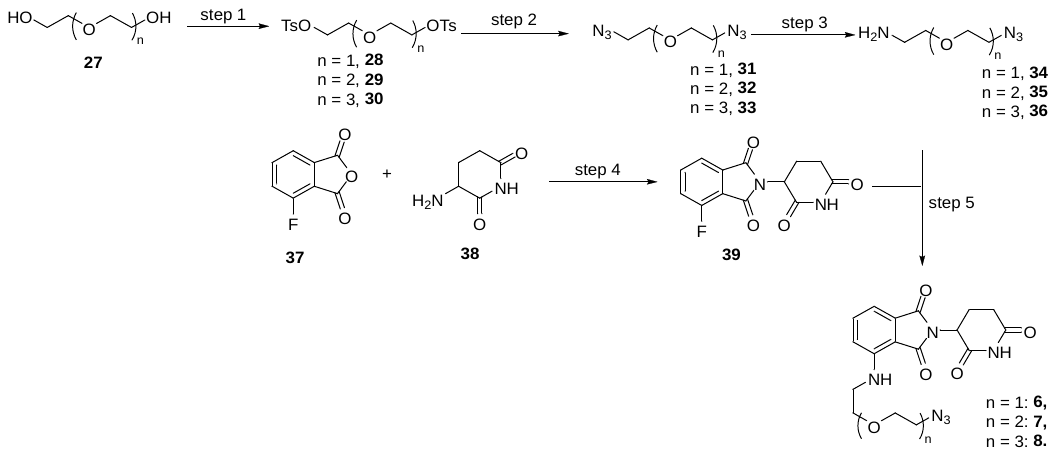

**1.1.2 General synthetic procedure for VHL ligand conjugates with Linkers 9-12.**

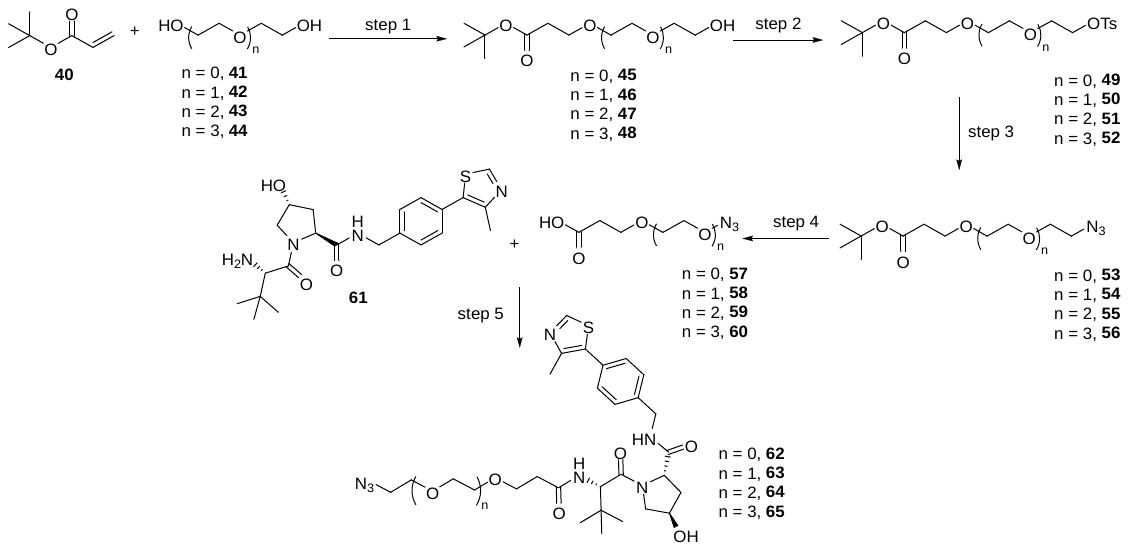

Step 1: Compound 40 (128 mg, 1.0 mmol) and the corresponding dihydroxy compounds (41–44, 5.0 mmol) were dissolved in dichloromethane (DCM, 10 mL) within a dry 25 mL round-bottom flask at ambient temperature. The reaction mixture was cooled to 0 °C under magnetic stirring. Triton B (0.05 mmol) was subsequently added. The reaction was then allowed to warm to room temperature. After stirring for 6 h, reaction completion was confirmed by LCMS analysis. The solvent was removed under reduced pressure. The crude product was purified by silica gel column chromatography to afford compounds 45–48 as colorless oils in yields ranging from 63% to 72%.

**1.1.3 Procedure for synthesis of intermediate 19**

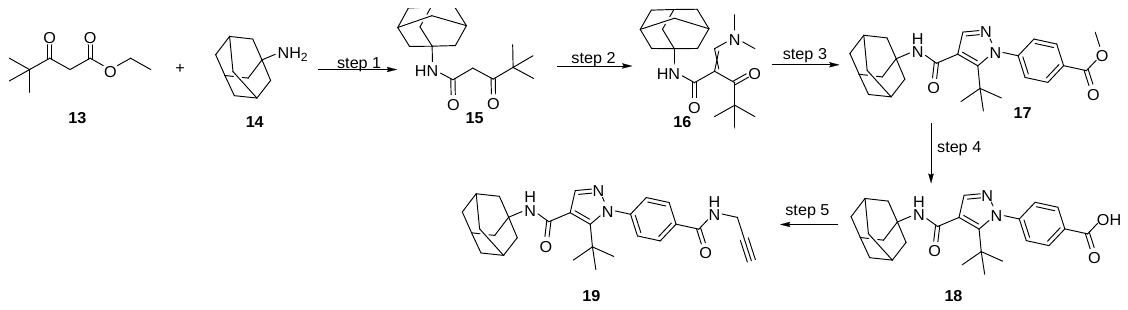
**N-(adamantan-1-yl)-4,4-dimethyl-3-oxopentanamide (15)**

Compound 13 (344 mg, 2 mmol) and compound 14 (302 mg, 2 mmol) were dissolved in xylene (10 mL) in a sealed tube. The reaction mixture was then placed under a nitrogen atmosphere and heated at 150 °C for 6 h. Thin-layer chromatography (TLC) indicated the complete consumption of starting materials. The crude product was purified by silica gel column chromatography to afford 487 mg of an off-white solid in 88% yield.

**1.1.4 General procedure for Click Reaction**

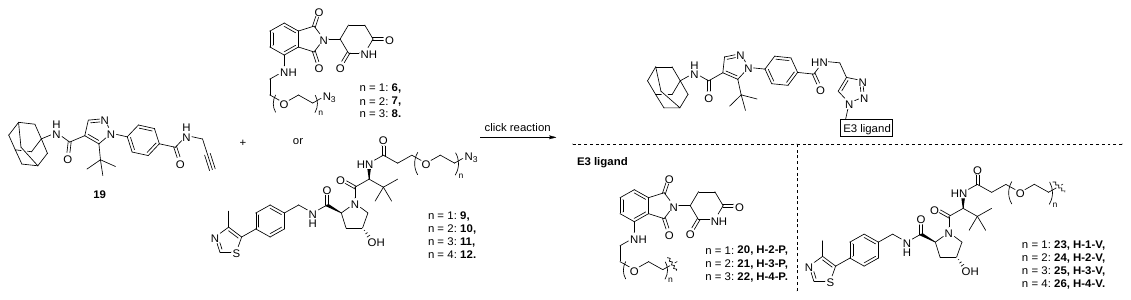
 Compound 19 (10 mg, 0.022 mmol, 1 equiv) and the corresponding E3 ligand (0.022 mmol, 1 equiv), synthesized as described in Sections 1.1.1 and 1.1.2, were dissolved in DMF (2 mL). A 1 M aqueous solution of CuSO₄ and a 1 M aqueous solution of sodium ascorbate (VicNa) were prepared. Subsequently, 0.05 equiv of the CuSO₄ solution and 0.10 equiv of the VicNa solution were added. The reaction mixture was stirred at ambient temperature for 12 h. Reaction completion was confirmed by LCMS analysis. The crude product was purified by silica gel column chromatography, affording the final compound as a white solid for the VH032-based derivative and as a yellow solid for the pomalidomide-based derivative in approximately 50% yield.

Compounds 20–26 were synthesized using the general procedure for Click Reaction.

**1.1.5 NMR and Mass Spectrometry Data for Compounds 20–26**

**N-(adamantan-1-yl)-5-(tert-butyl)-1-(4-(((1-(2-(2-((2-(2,6-dioxopiperidin-3-yl)-1,3-dioxoisoindolin-4-yl)amino)ethoxy)ethyl)-1H-1,2,3-triazol-4-yl)methyl)carbamoyl)phenyl)-1H-pyrazole-4-carboxamide (20)**

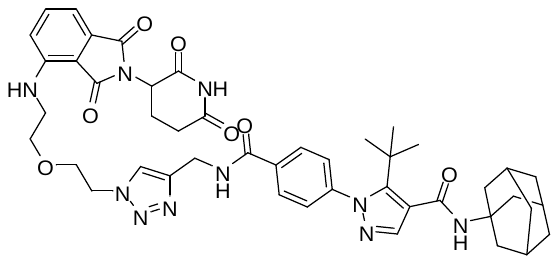

^1^H NMR (300 MHz, DMSO-*d*_6_) δ 11.10 (s, 1H), 9.27 – 9.12 (m, 1H), 8.07 – 7.91 (m, 3H), 7.72 (s, 1H), 7.63 – 7.50 (m, 2H), 7.44 (d, *J* = 8.1 Hz, 2H), 7.08 (dd, *J* = 19.6, 7.8 Hz, 2H), 6.60 (t, *J* = 5.8 Hz, 1H), 5.08 (dd, *J* = 12.5, 5.2 Hz, 1H), 4.63 – 4.44 (m, 4H), 3.86 (t, *J* = 5.3 Hz, 2H), 3.61 (d, *J* = 5.4 Hz, 2H), 3.52 – 3.38 (m, 2H), 2.96 – 2.82 (m, 1H), 2.65 – 2.54 (m, 2H), 2.11 – 1.95 (m, 11H), 1.65 (s, 5H), 1.19 (s, 9H).

^13^C NMR (75 MHz, DMSO) δ 172.78, 170.07, 168.92, 167.27, 165.23, 164.83, 148.99, 146.35, 144.69, 144.58, 138.83, 136.21, 134.72, 132.07, 128.01, 127.93, 123.46, 119.41, 117.41, 110.69, 109.25, 68.86, 68.70, 51.25, 49.23, 48.55, 41.54, 40.75, 36.11, 34.90, 33.03, 30.98, 30.45, 28.82, 22.14.

HRMS (m/z):

calculated for C_45_H_53_N_10_O_7_ [M+H]^+^: 845.4099; found: 845.4108

calculated for C_45_H_52_N_10_O_7_Na [M+Na]^+^: 867.3918; found: 867.3950.

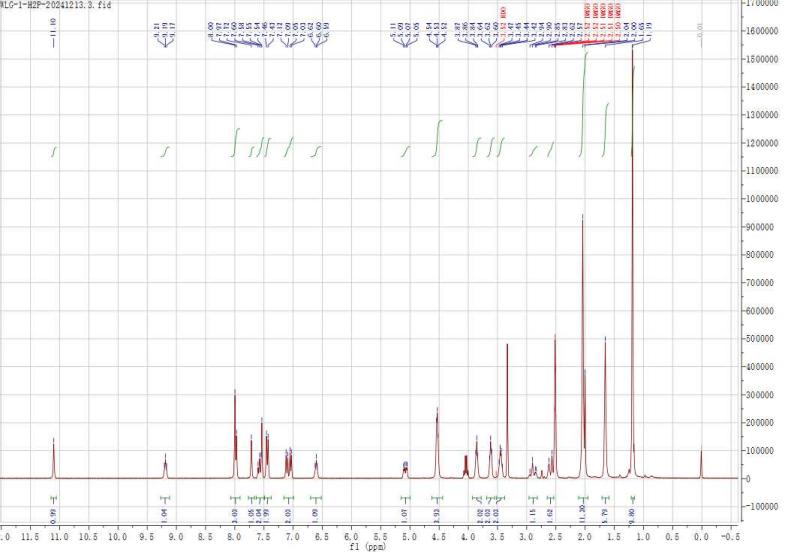

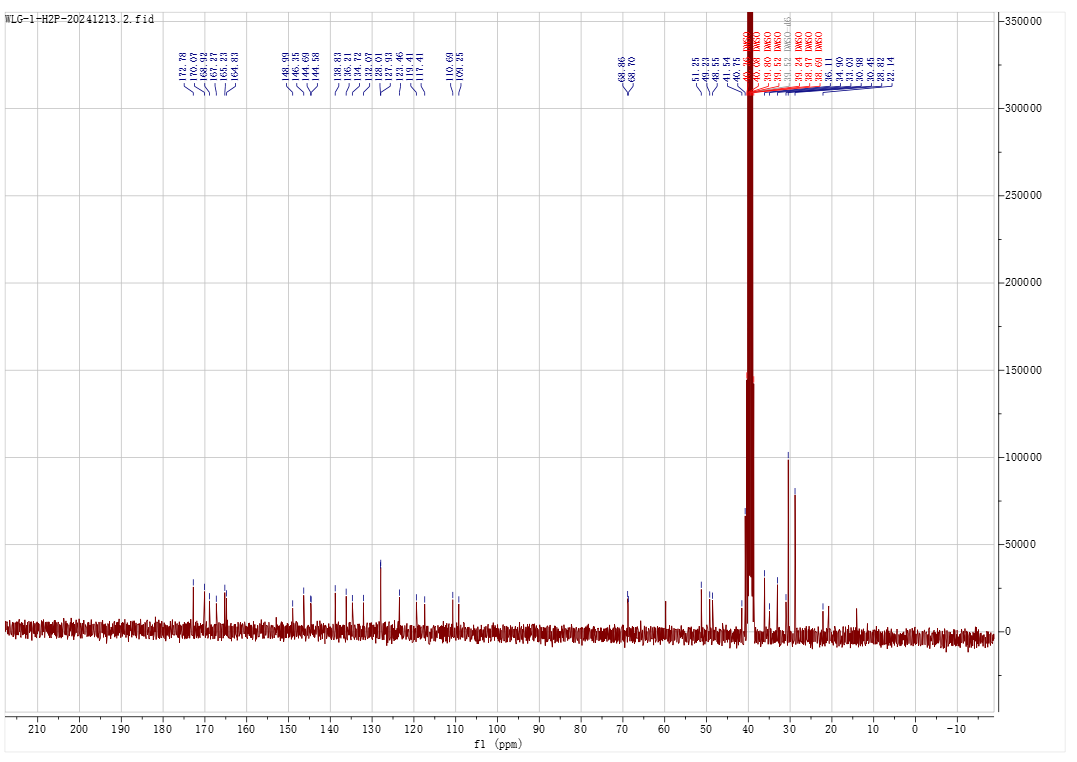

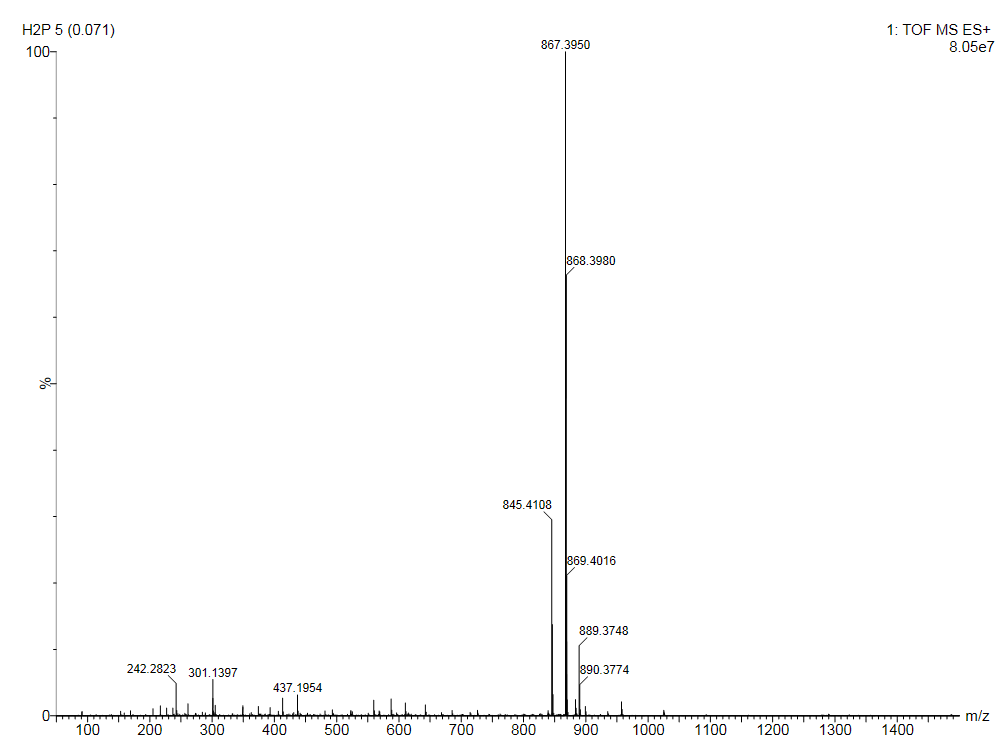

**N-(adamantan-1-yl)-5-(tert-butyl)-1-(4-(((1-(2-(2-(2-((2-(2,6-dioxopiperidin-3-yl)-1,3-dioxoisoindolin-4-yl)amino)ethoxy)ethoxy)ethyl)-1H-1,2,3-triazol-4-yl)methyl)carbamoyl)phenyl)-1H-pyrazole-4-carboxamide (21)**

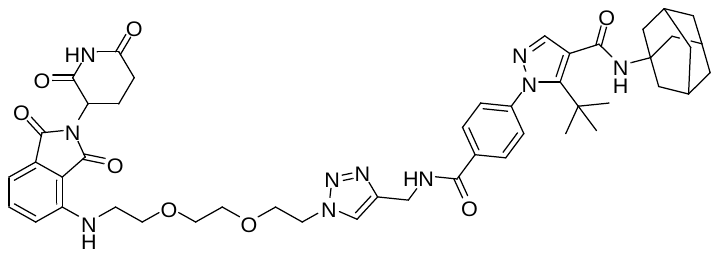

^1^H NMR (300 MHz, DMSO-*d*_6_) δ 11.08 (s, 1H), 9.20 (t, *J* = 5.7 Hz, 1H), 8.03 – 7.93 (m, 3H), 7.70 (s, 1H), 7.63 – 7.49 (m, 2H), 7.48 – 7.39 (m, 2H), 7.12 (d, *J* = 8.6 Hz, 1H), 7.03 (d, *J* = 7.1 Hz, 1H), 6.64 – 6.54 (m, 1H), 5.05 (dd, *J* = 12.9, 5.2 Hz, 1H), 4.56 – 4.43 (m, 4H), 3.81 (t, *J* = 5.2 Hz, 2H), 3.61 – 3.49 (m, 6H), 3.44 (t, *J* = 5.3 Hz, 2H), 2.96 – 2.78 (m, 1H), 2.64 – 2.51 (m, 2H), 2.06 – 1.96 (m, 11H), 1.64 (s, 6H), 1.18 (s, 9H).

^13^C NMR (75 MHz, DMSO) δ 172.78, 170.07, 168.95, 167.27, 165.22, 164.83, 149.00, 146.38, 144.73, 144.60, 138.83, 136.22, 134.70, 132.07, 128.03, 127.92, 123.37, 119.41, 117.43, 110.68, 109.23, 69.62, 69.56, 68.84, 68.79, 51.26, 49.30, 48.54, 41.64, 40.76, 36.11, 34.92, 33.03, 30.97, 30.45, 28.83, 22.12.

HRMS (m/z):

calculated for C_47_H_57_N_10_O_8_ [M+H]^+^: 889.4361; found: 889.4399

calculated for C_47_H_56_N_10_O_8_Na [M+Na]^+^: 911.4180; found: 911.4188.

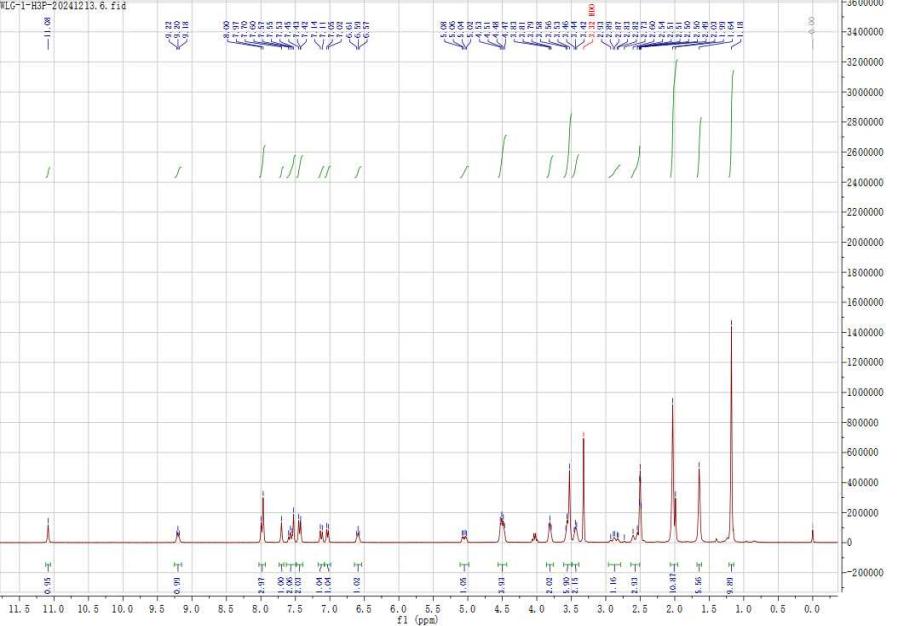

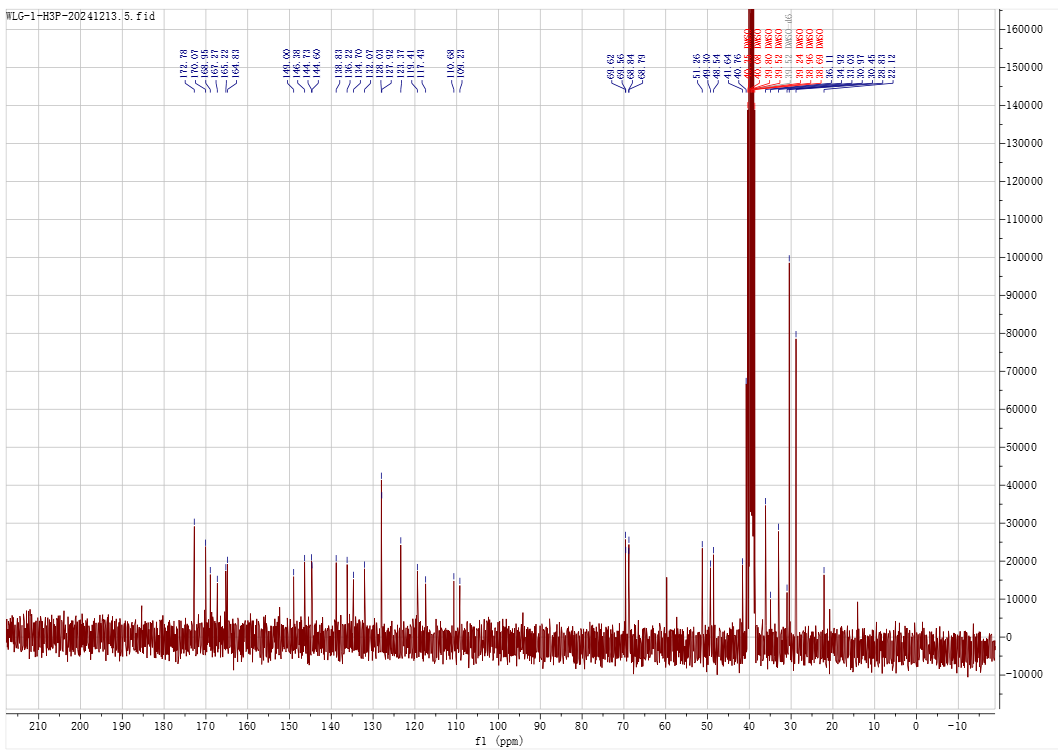

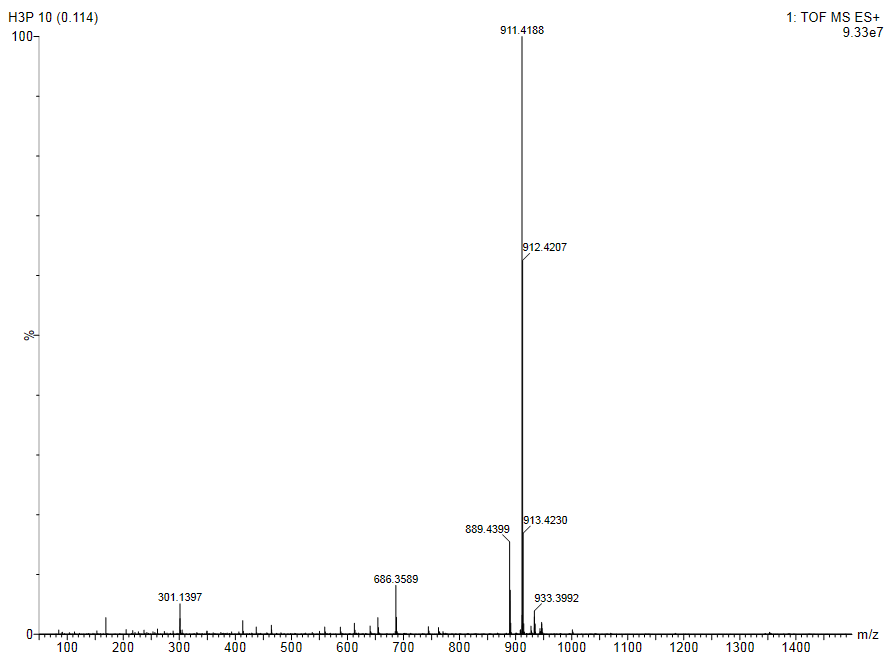

**N-(adamantan-1-yl)-5-(tert-butyl)-1-(4-(((1-(2-(2-(2-(2-((2-(2,6-dioxopiperidin-3-yl)-1,3-dioxoisoindolin-4-yl)amino)ethoxy)ethoxy)ethoxy)ethyl)-1H-1,2,3-triazol-4-yl)methyl)carbamoyl)phenyl)-1H-pyrazole-4-carboxamide (22)**

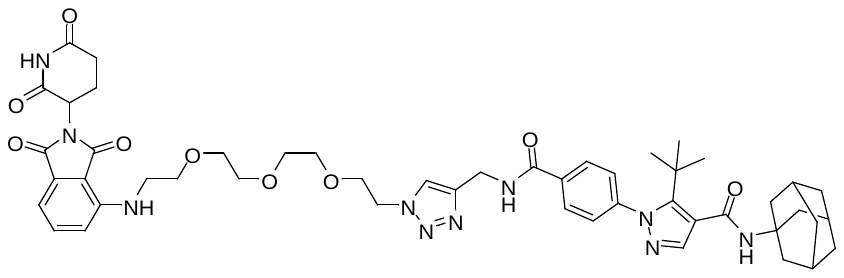

^1^H NMR (300 MHz, DMSO) δ 11.09 (s, 1H), 9.20 (t, *J* = 5.8 Hz, 1H), 8.10 – 7.89 (m, 3H), 7.70 (s, 1H), 7.63 – 7.50 (m, 2H), 7.44 (d, *J* = 8.1 Hz, 2H), 7.13 (d, *J* = 8.6 Hz, 1H), 7.03 (d, *J* = 7.1 Hz, 1H), 6.59 (t, *J* = 5.8 Hz, 1H), 5.05 (dd, *J* = 12.8, 5.3 Hz, 1H), 4.57 – 4.43 (m, 4H), 3.79 (t, *J* = 5.3 Hz, 2H), 3.62 – 3.42 (m, 12H), 2.96 – 2.79 (m, 1H), 2.64 – 2.50 (m, 2H), 2.03 (s, 10H), 1.64 (s, 6H), 1.18 (s, 9H).

^13^C NMR (75 MHz, DMSO) δ 172.79, 170.06, 168.93, 167.28, 165.21, 164.83, 149.00, 146.39, 144.73, 144.61, 138.83, 136.21, 134.70, 132.08, 128.03, 127.93, 123.36, 119.42, 117.44, 110.66, 109.22, 69.74, 69.69, 69.53, 68.84, 68.72, 51.25, 49.27, 48.54, 41.67, 40.75, 36.10, 34.92, 33.04, 30.97, 30.45, 28.83, 22.13.

HRMS (m/z):

calculated for C_49_H_61_N_10_O_9_ [M+H]^+^: 933.4623; found: 933.4659

calculated for C_49_H_60_N_10_O_9_Na [M+Na]^+^: 955.4442; found: 955.4392.

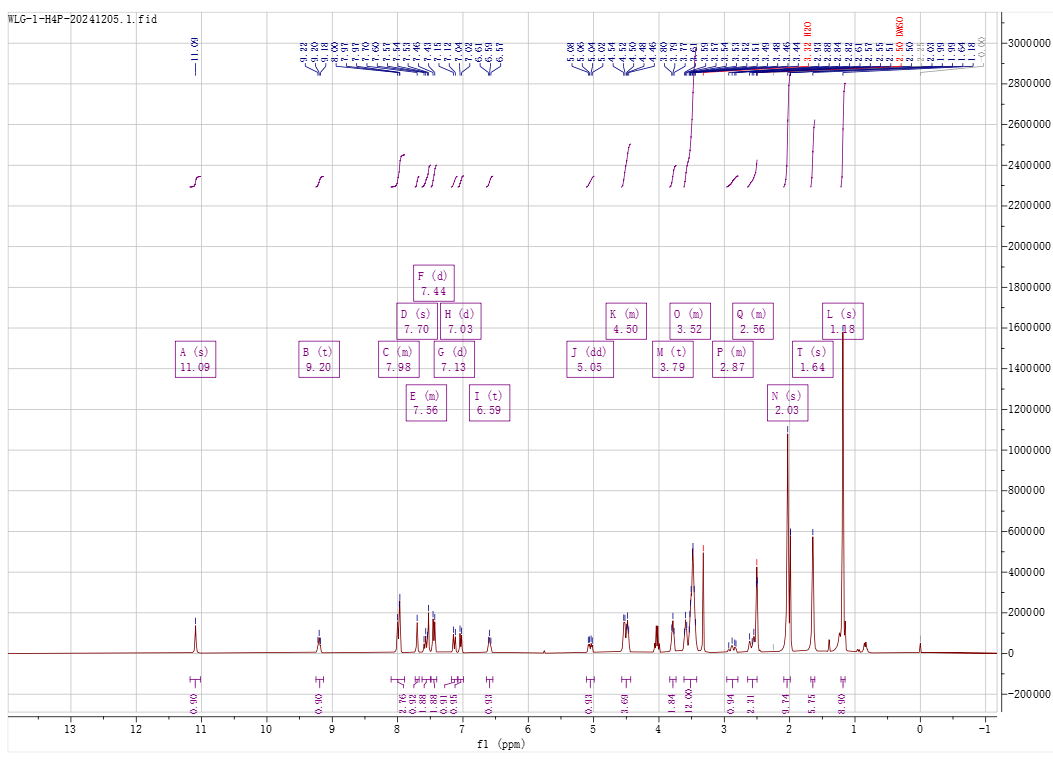

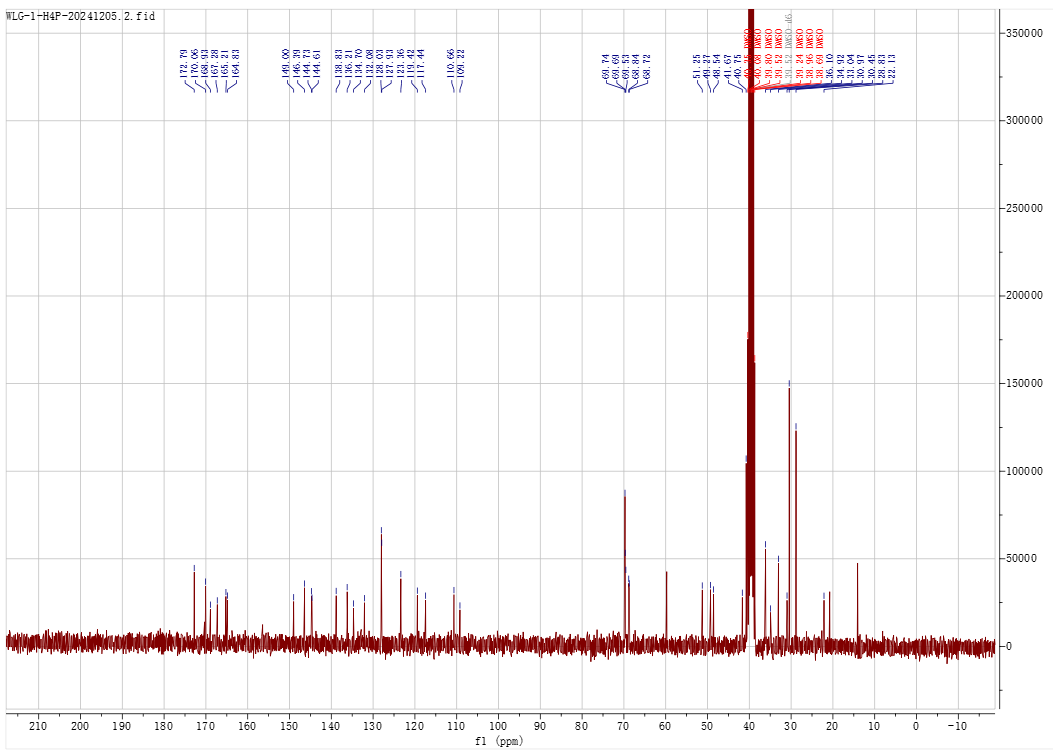

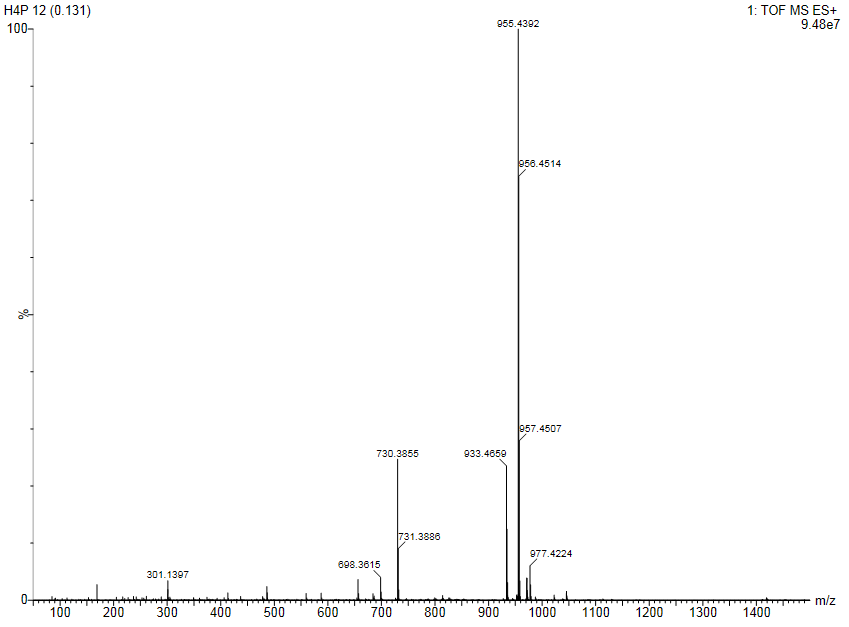

**N-(adamantan-1-yl)-5-(tert-butyl)-1-(4-(((1-(2-(3-(((S)-1-((2S,4R)-4-hydroxy-2-((4-(4-methylthiazol-5-yl)benzyl)carbamoyl)pyrrolidin-1-yl)-3,3-dimethyl-1-oxobutan-2-yl)amino)-3-oxopropoxy)ethyl)-1H-1,2,3-triazol-4-yl)methyl)carbamoyl)phenyl)-1H-pyrazole-4-carboxamide (23)**

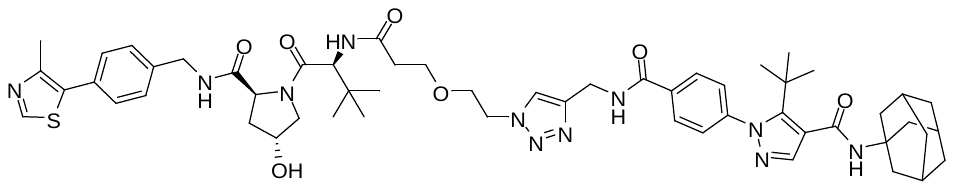

^1^H NMR (300 MHz, DMSO) δ 9.19 (t, *J* = 5.7 Hz, 1H), 8.98 (s, 1H), 8.57 (t, *J* = 6.0 Hz, 1H), 8.04 – 7.90 (m, 4H), 7.71 (s, 1H), 7.52 (s, 1H), 7.49 – 7.32 (m, 6H), 5.14 (t, *J* = 3.6 Hz, 1H), 4.58 – 4.17 (m, 9H), 3.84 – 3.53 (m, 6H), 2.55 (d, *J* = 6.5 Hz, 1H), 2.44 (d, *J* = 1.4 Hz, 3H), 2.36 (dt, *J* = 13.9, 6.2 Hz, 1H), 2.03 (s, 10H), 1.96 – 1.83 (m, 1H), 1.64 (s, 6H), 1.19 (s, 9H), 0.92 (s, 9H).

^13^C NMR (75 MHz, DMSO) δ 171.87, 169.78, 169.53, 165.23, 164.81, 151.39, 148.99, 147.69, 144.65, 144.59, 139.45, 138.81, 134.71, 131.13, 129.62, 128.60, 128.00, 127.93, 127.39, 123.37, 119.40, 68.84, 68.40, 66.74, 58.68, 56.35, 51.25, 49.23, 41.62, 40.75, 37.93, 36.09, 35.42, 35.27, 35.13, 34.91, 33.02, 30.45, 28.81, 26.31, 15.91.

HRMS (m/z):

calculated for C_55_H_72_N_11_O_7_S [M+H]^+^: 1030.5337; found: 1030.5295

calculated forC_55_H_71_N_11_O_7_SNa [M+Na]^+^: 1052.5156; found: 1052.5198.

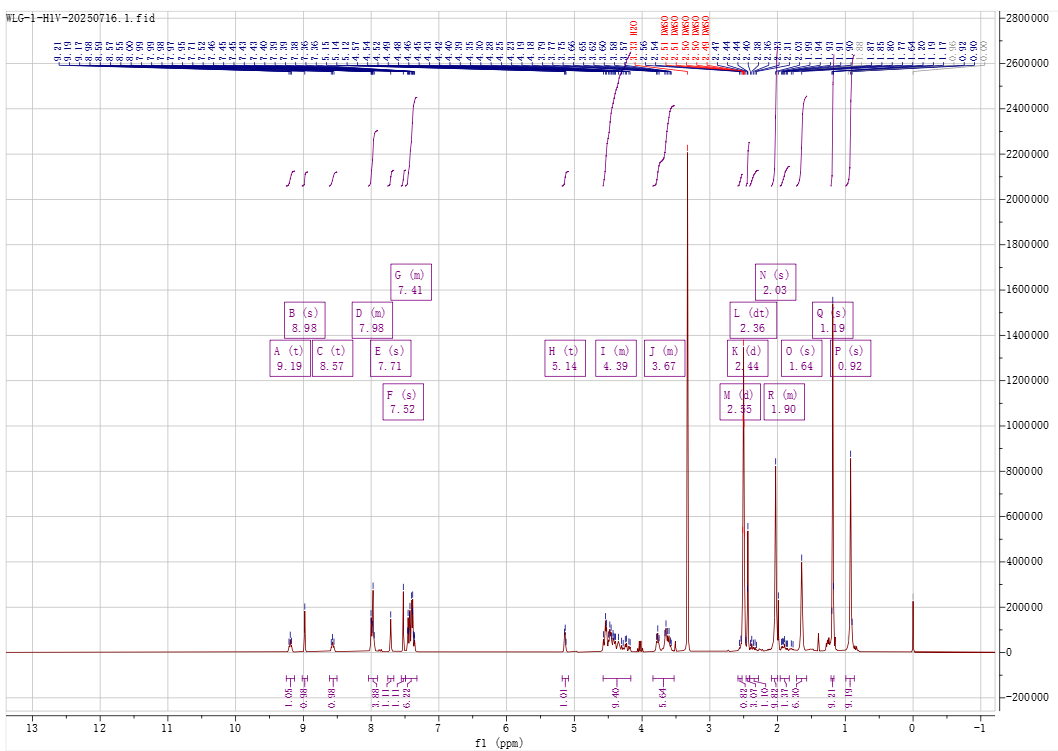

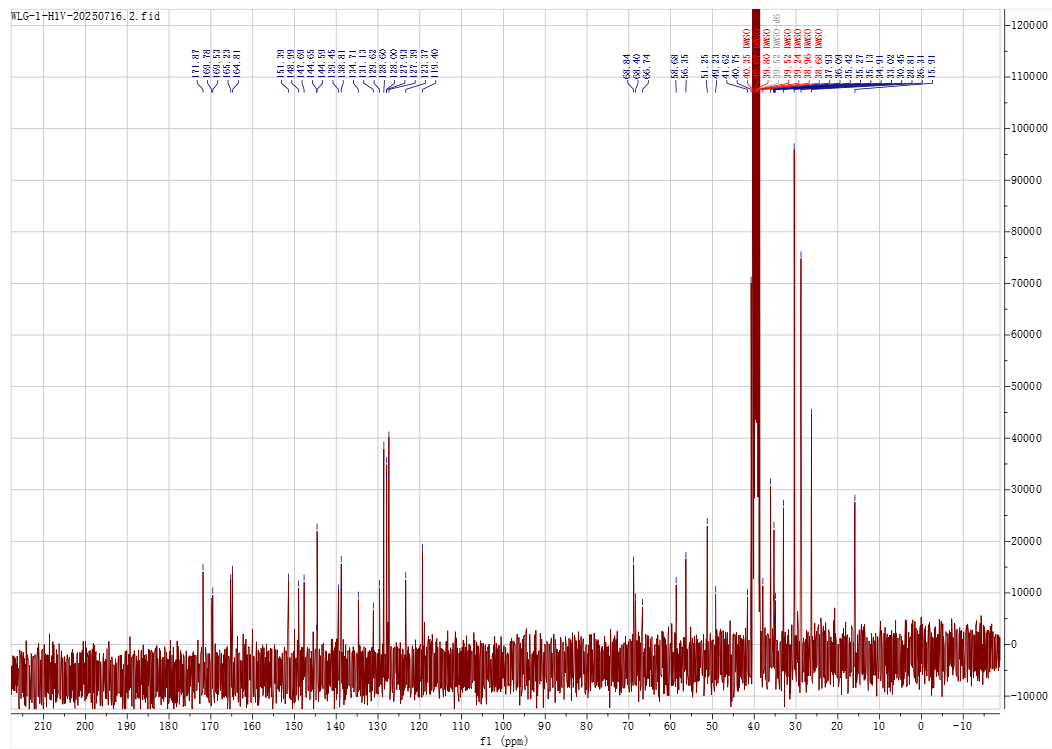

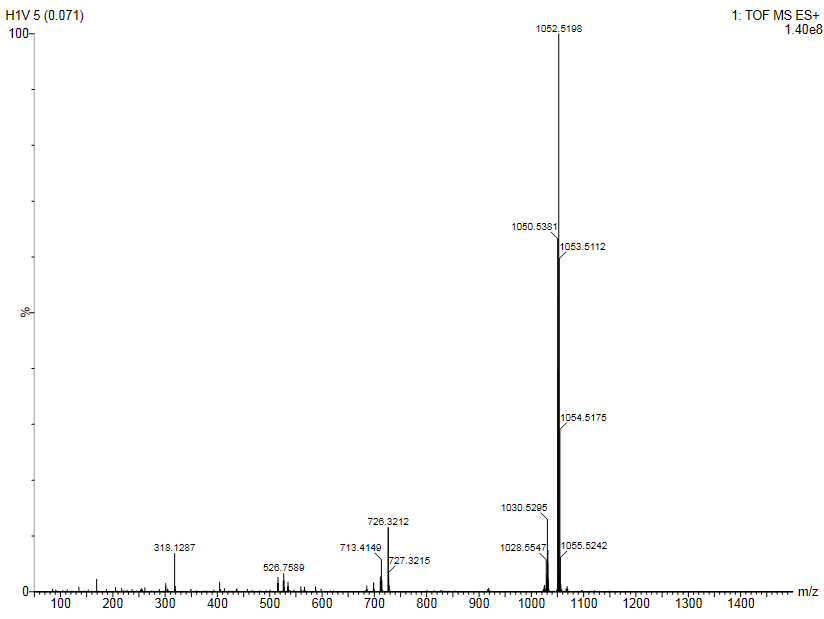

**N-(adamantan-1-yl)-5-(tert-butyl)-1-(4-(((1-(2-(2-(3-(((S)-1-((2S,4R)-4-hydroxy-2-((4-(4-methylthiazol-5-yl)benzyl)carbamoyl)pyrrolidin-1-yl)-3,3-dimethyl-1-oxobutan-2-yl)amino)-3-oxopropoxy)ethoxy)ethyl)-1H-1,2,3-triazol-4-yl)methyl)carbamoyl)phenyl)-1H-pyrazole-4-carboxamide (24)**

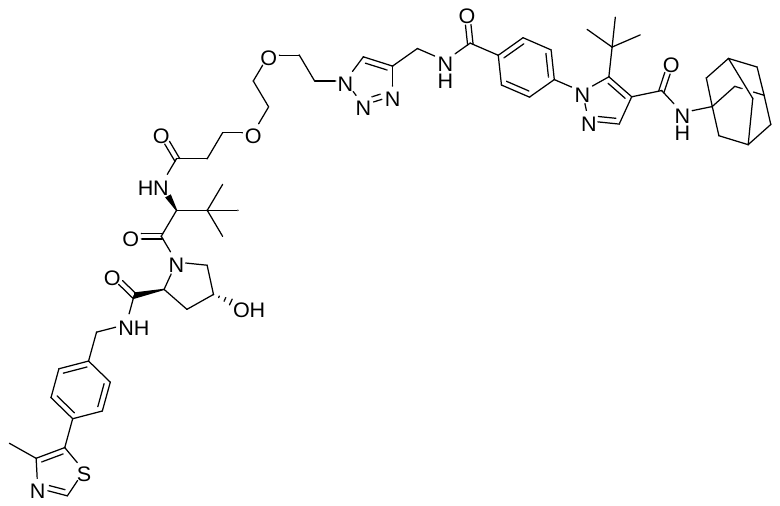

^1^H NMR (300 MHz, DMSO) δ 9.21 (s, 1H), 8.98 (s, 1H), 8.56 (s, 1H), 8.05 – 7.87 (m, 4H), 7.71 (s, 1H), 7.53 (s, 1H), 7.50 – 7.35 (m, 6H), 5.19 – 5.08 (m, 1H), 4.59 – 4.32 (m, 8H), 4.29 – 4.17 (m, 1H), 3.79 (t, *J* = 5.2 Hz, 2H), 3.68 – 3.40 (m, 8H), 2.56 (s, 1H), 2.44 (s, 3H), 2.41 – 2.27 (m, 1H), 2.03 (s, 10H), 1.90 (s, 1H), 1.64 (s, 6H), 1.19 (s, 9H), 0.93 (s, 9H).

^13^C NMR (75 MHz, DMSO) δ 171.90, 169.94, 169.52, 165.25, 164.84, 151.44, 149.00, 147.71, 144.72, 144.60, 139.48, 138.82, 134.72, 131.14, 129.63, 128.63, 128.03, 127.95, 127.41, 123.38, 119.41, 69.43, 69.36, 68.86, 68.75, 66.90, 58.70, 56.36, 56.29, 51.25, 49.26, 41.64, 40.75, 37.96, 36.10, 35.62, 35.35, 34.93, 33.04, 30.46, 28.82, 26.31, 15.93.

HRMS (m/z):

calculated for C_57_H_76_N_11_O_8_S [M+H]^+^: 1074.5599; found: 1074.5560

calculated forC_57_H_75_N_11_O_8_SNa [M+Na]^+^: 1096.5418; found: 1096.5460.

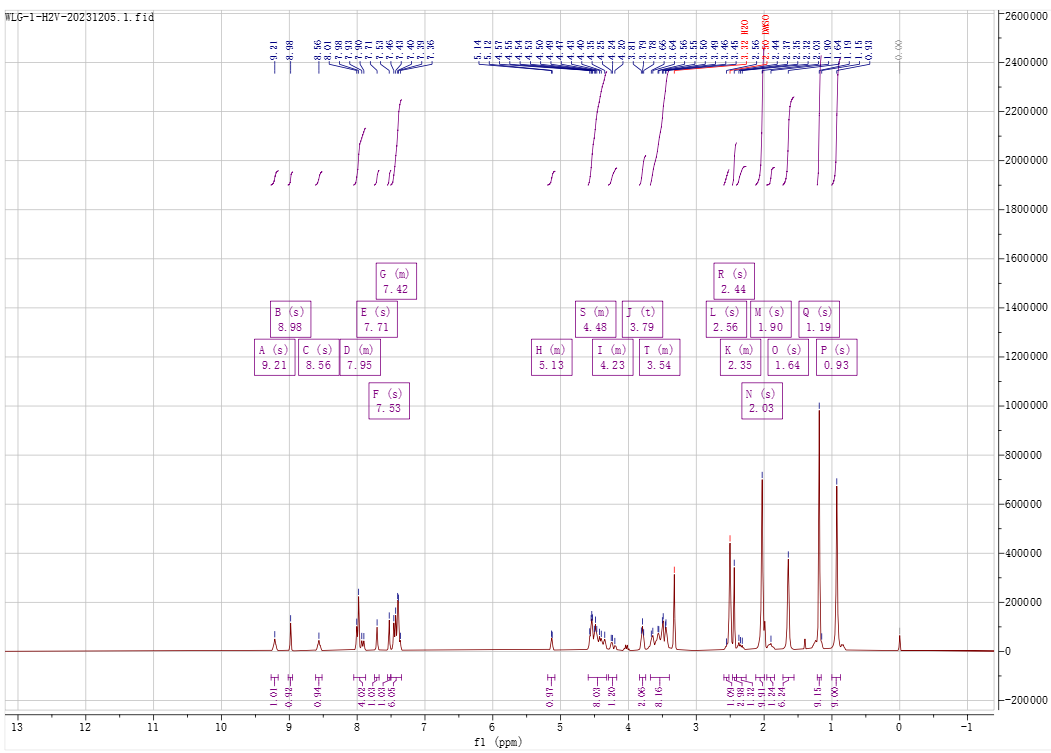

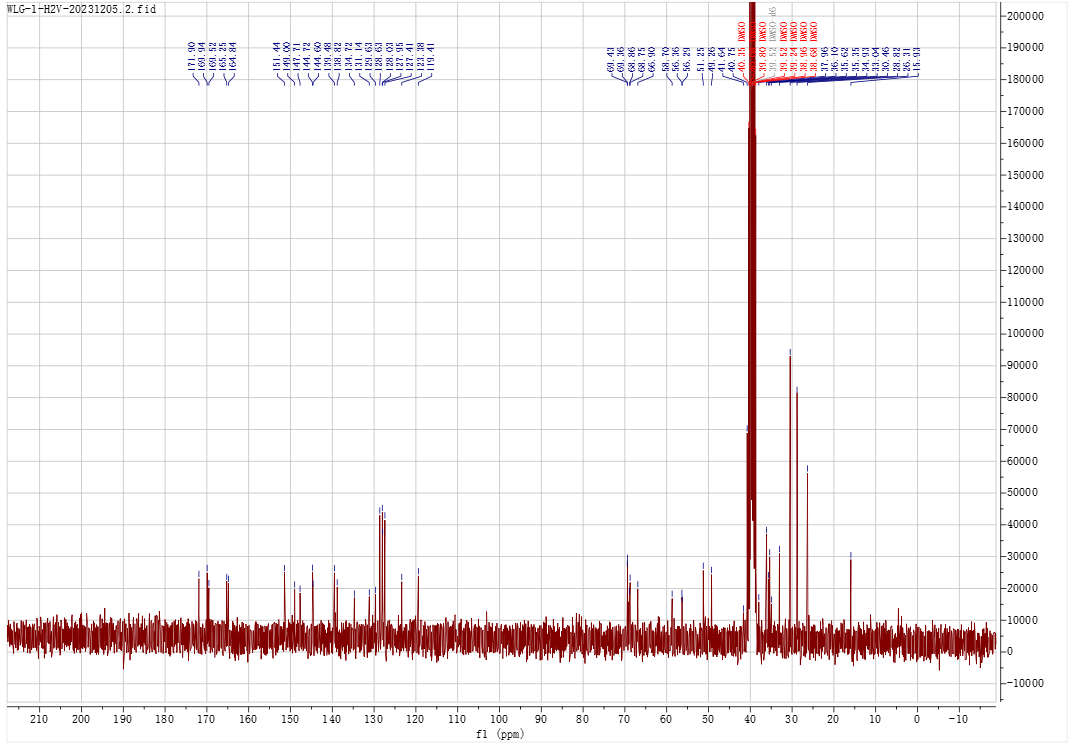

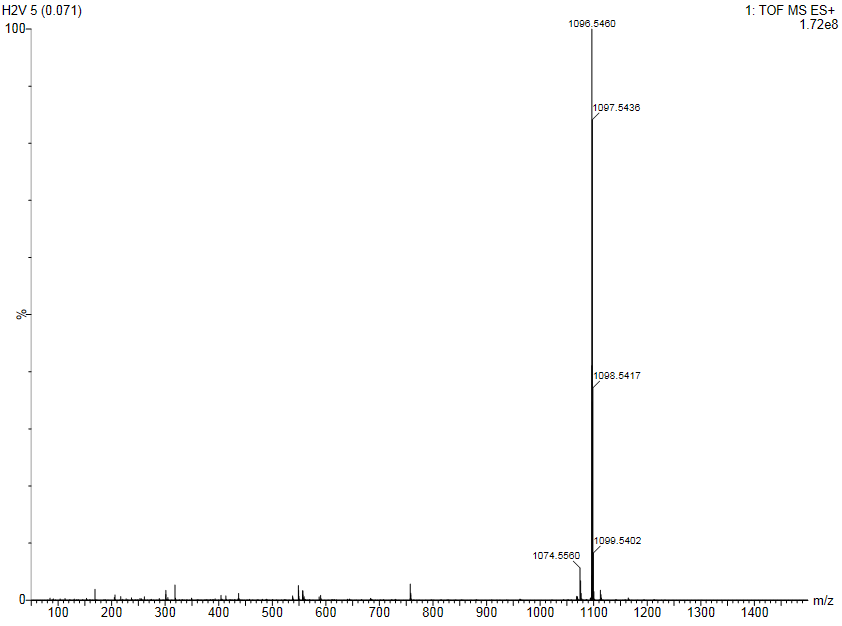

**N-(adamantan-1-yl)-5-(tert-butyl)-1-(4-(((1-((S)-14-((2S,4R)-4-hydroxy-2-((4-(4-methylthiazol-5-yl)benzyl)carbamoyl)pyrrolidine-1-carbonyl)-15,15-dimethyl-12-oxo-3,6,9-trioxa-13-azahexadecyl)-1H-1,2,3-triazol-4-yl)methyl)carbamoyl)phenyl)-1H-pyrazole-4-carboxamide (25)**

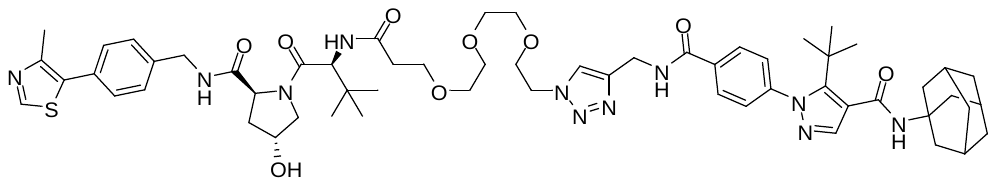

^1^H NMR (300 MHz, DMSO) δ 9.26 – 9.15 (m, 1H), 8.98 (s, 1H), 8.56 (t, *J* = 6.0 Hz, 1H), 8.04 – 7.88 (m, 4H), 7.71 (s, 1H), 7.53 (s, 1H), 7.49 – 7.34 (m, 6H), 5.13 (d, *J* = 3.4 Hz, 1H), 4.60 – 4.31 (m, 8H), 4.22 (dd, *J* = 16.0, 5.4 Hz, 1H), 3.80 (t, *J* = 5.3 Hz, 2H), 3.71 – 3.40 (m, 12H), 2.59 – 2.52 (m, 1H), 2.41 – 2.28 (m, 1H), 2.01 (s, 10H), 1.96 – 1.83 (m, 1H), 1.64 (s, 6H), 1.19 (s, 9H), 0.93 (s, 9H).

^13^C NMR (75 MHz, DMSO) δ 171.90, 169.92, 169.52, 165.21, 164.84, 151.44, 149.00, 147.70, 144.73, 144.60, 139.49, 138.83, 134.71, 131.15, 129.63, 128.63, 128.03, 127.93, 127.41, 123.37, 119.41, 69.67, 69.63, 69.49, 69.45, 68.85, 68.73, 66.92, 58.69, 56.35, 56.28, 51.26, 49.27, 41.63, 40.75, 37.93, 36.10, 35.63, 35.35, 34.93, 33.04, 30.46, 28.82, 26.32, 15.93.

HRMS (m/z):

calculated for C_59_H_80_N_11_O_9_S [M+H]^+^: 1118.5861; found: 1118.5835

calculated forC_59_H_79_N_11_O_9_SNa [M+Na]^+^: 1140.5681; found: 1140.5750.

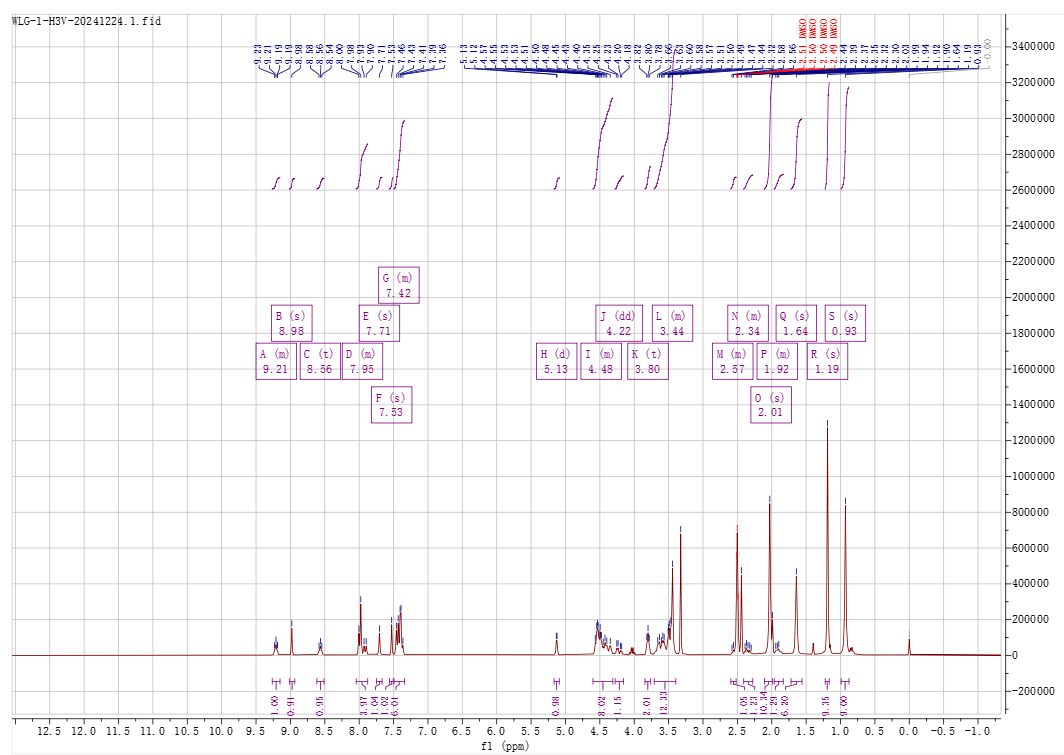

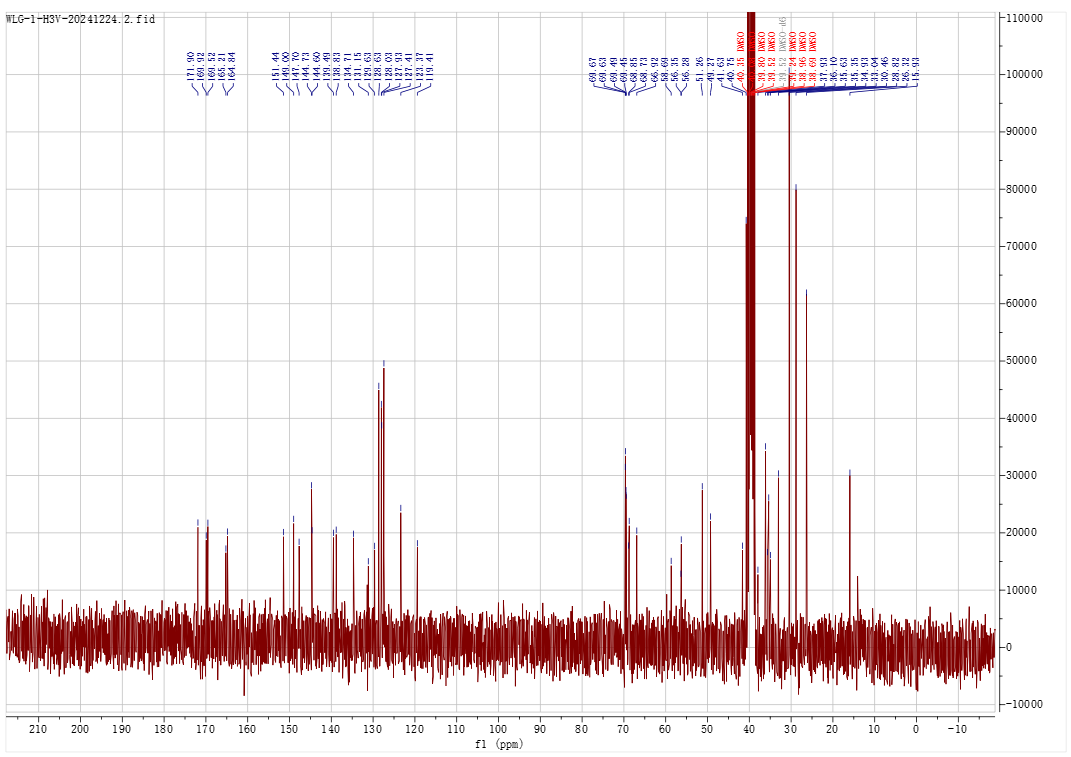

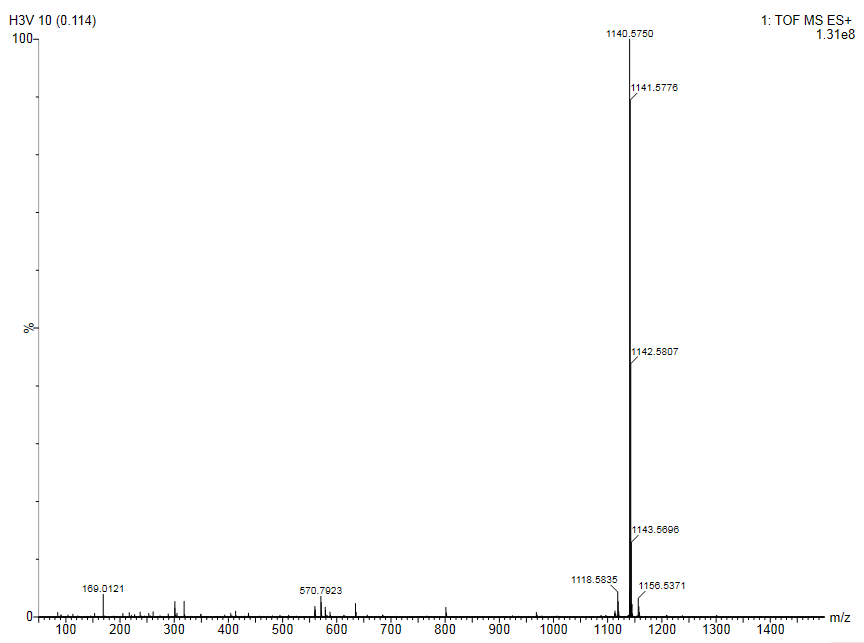

**N-(adamantan-1-yl)-5-(tert-butyl)-1-(4-(((1-((S)-17-((2S,4R)-4-hydroxy-2-((4-(4-methylthiazol-5-yl)benzyl)carbamoyl)pyrrolidine-1-carbonyl)-18,18-dimethyl-15-oxo-3,6,9,12-tetraoxa-16-azanonadecyl)-1H-1,2,3-triazol-4-yl)methyl)carbamoyl)phenyl)-1H-pyrazole-4-carboxamide (26)**

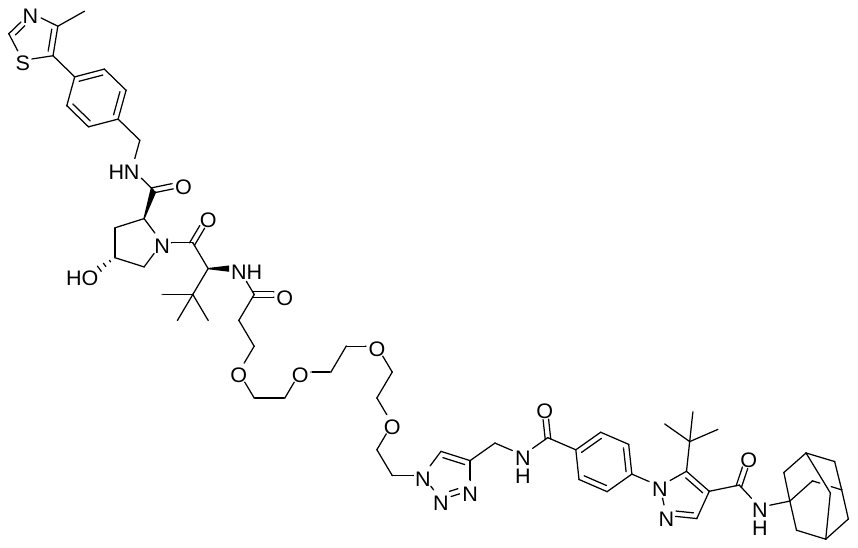

^1^H NMR (300 MHz, DMSO) δ 9.21 (s, 1H), 8.98 (s, 1H), 8.55 (s, 1H), 8.02 – 7.88 (m, 4H), 7.71 (s, 1H), 7.53 (s, 1H), 7.49 – 7.34 (m, 6H), 5.13 (d, *J* = 3.3 Hz, 1H), 4.60 – 4.31 (m, 8H), 4.22 (dd, *J* = 15.5, 5.2 Hz, 1H), 3.80 (t, *J* = 5.3 Hz, 2H), 3.72 – 3.37 (m, 16H), 2.60 – 2.52 (m, 1H), 2.44 (s, 3H), 2.40 – 2.26 (m, 1H), 2.03 (s, 10H), 1.96 – 1.82 (m, 1H), 1.64 (s, 6H), 1.19 (s, 9H), 0.93 (s, 9H).

^13^C NMR (75 MHz, DMSO) δ 171.91, 169.91, 169.52, 165.21, 164.84, 151.44, 148.99, 147.71, 144.72, 144.61, 139.49, 138.83, 134.71, 131.14, 129.63, 128.63, 128.03, 127.93, 127.41, 123.37, 119.41, 69.73, 69.67, 69.62, 69.51, 69.44, 68.86, 68.74, 66.93, 58.70, 56.36, 56.27, 51.26, 49.27, 41.64, 40.75, 37.93, 36.11, 35.64, 35.35, 34.93, 33.04, 30.46, 28.82, 26.32, 15.93.

HRMS (m/z):

calculated for C_61_H_84_N_11_O_10_S [M+H]^+^: 1162.6123; found: 1162.6167

calculated forC_61_H_83_N_11_O_10_SNa [M+Na]^+^: 1184.5943; found: 1184.5989.

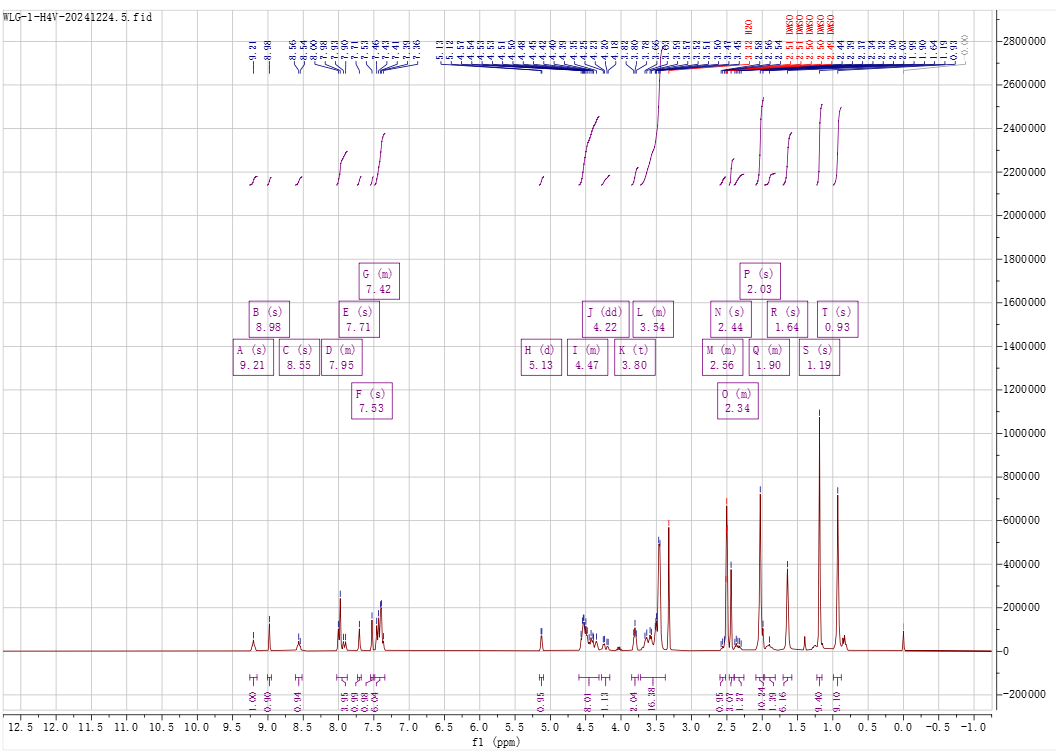

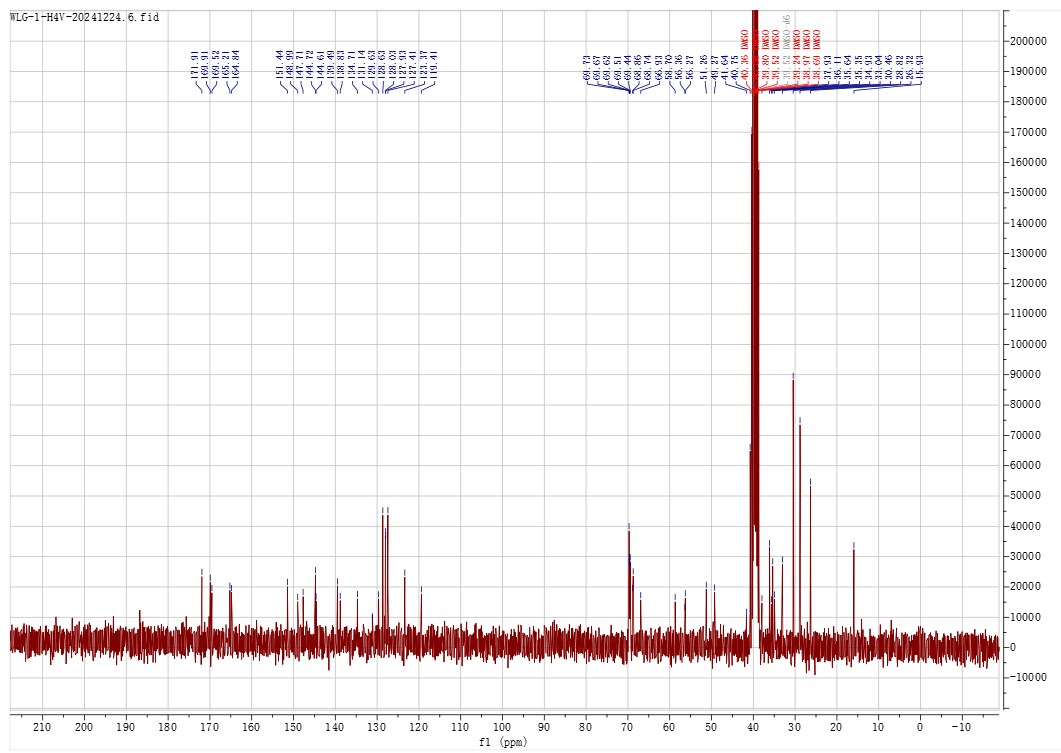

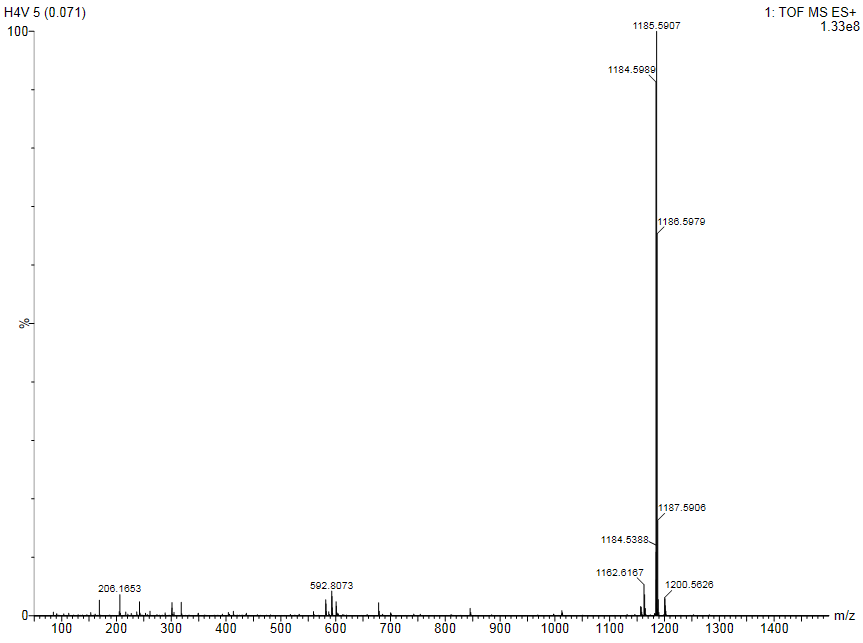

**2 Cell line and culture**

The human biphenotypic B myelomonocytic leukemia cell line MV4-11 was obtained from Procell system (CL-0572). Cells were cultured in Iscove's Modified Dulbecco's Medium (IMDM) (Gibco, C12440500BT) supplemented with 10% fetal bovine serum (FBS) (CELLiGENT, China, Cat#:CG1126B) and 1% penicillin-streptomycin (P/S) (AbbKine, USA, Cat#: BMC1030). Cells were maintained in sterile culture flasks at 37°C in a humidified incubator with 5% CO₂. The medium was refreshed every 48 hours to maintain optimal nutrient conditions. Cell density was routinely monitored and maintained between 1×10⁵ and 1×10⁶ cells/mL by subculturing or dilution with fresh medium. For cell treatment, MV4-11 cells were seeded in 6-well plates at a density of 5 x 10^6^ cells per well. After 24-hour incubation, cells were treated with the following compounds, drug or vehicle controls such as H-1-P, H-2-P, H-3-P, H-1-V, H-2-V, H-3-V, H-4-V, pomalidomide (bidepharm, BD235626), VH032 (bidepharm, BD00758812), bortezomib (bidepharm, BD106087), carfilzomib (bidepharm, BD292920), compound 18 and DMSO, for specified durations. After treatment, cells were lysed and subjected to Western blot analysis to determine 11β-HSD1 protein expression.

**7 Molecular docking and dynamics stimulation**

First, the AZD8329-11β-HSD1 complex (PDB ID: 4P38) and the VH032-VHL complex (PDB ID: 8BDI) were downloaded using Schrödinger Suite 2024-2. The proteins were preprocessed using the Protein Preparation Wizard, which involved removing extraneous co-crystallized components, assigning bond orders, retaining only the monomeric protein and its cognate ligand, and modeling missing side chains. Grid files were subsequently generated around the ligand binding sites using default parameters. The molecular structures of H-1-V and H-3-V were sketched. An initial ternary complex system was constructed via molecular docking and structural alignment. Molecular dynamics (MD) simulations were performed using GROMACS (version 2025-2), employing the amber14sb_parmbsc1 force field for proteins and the GAFF force field for ligands. Ligand partial atomic charges were derived using Multiwfn and the RESP charge calculation script^2,3^, while the complete ligand topology was generated using AmberTools antechamber and converted to GROMACS format via acpype.py. Pre-equilibration and NPT simulations with position restraints were conducted to eliminate unfavorable interactions and generate initial atomic velocities. Following restraint removal, production MD simulations were performed to sample the conformational space. After 2 μs of cumulative simulation time, the binding free energy (ΔG) was calculated using gmx_MMPBSA^4^ according to the equation: ΔG = 〈G_Complex_〉 - 〈G_11β-HSD1_〉 - 〈G_VHL_〉 - 〈G_ligand_〉. This involved computing per-frame energy terms for the Ligand, 11β-HSD1, VHL, and the Complex. Trajectory analysis was performed using MDAnalysis Python scripts to calculate root-mean-square deviation (RMSD), root-mean-square fluctuation (RMSF), hydrogen bond occupancy, and principal component analysis (PCA). A free energy landscape (FEL) was constructed based on PCA projections. Conformational energies were then computed using the MMGBSA method (igb=8 model) indexed against the FEL, enabling the generation of a composite energy distribution plot. At the same time, we have also disclosed our analysis code, please see <https://github.com/wangl-g/trajectory_analysis/blob/main/trajectory_analysis>.

**Raw Data**

**1 Western blot**

Figure 3A 11β-HSD1

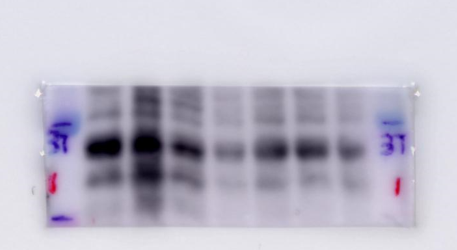

Figure 3A Tubulin

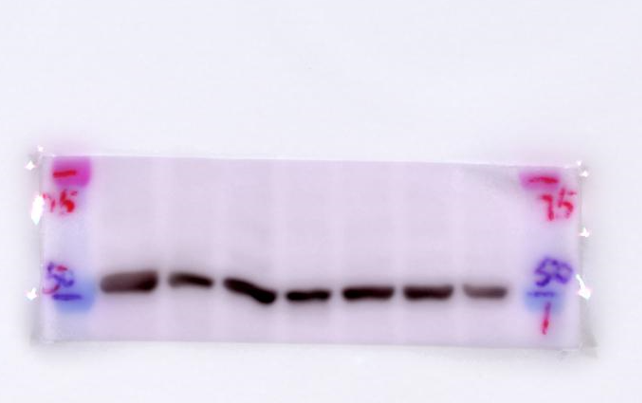

Figure	3B	11β-HSD1

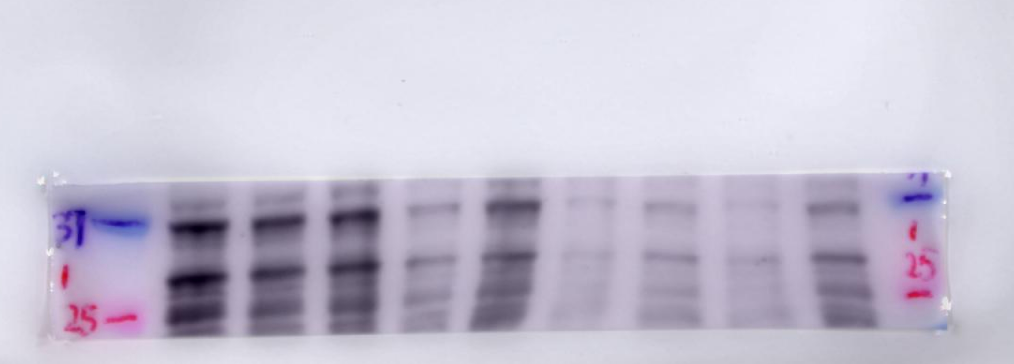

Figure	3B	Tubulin

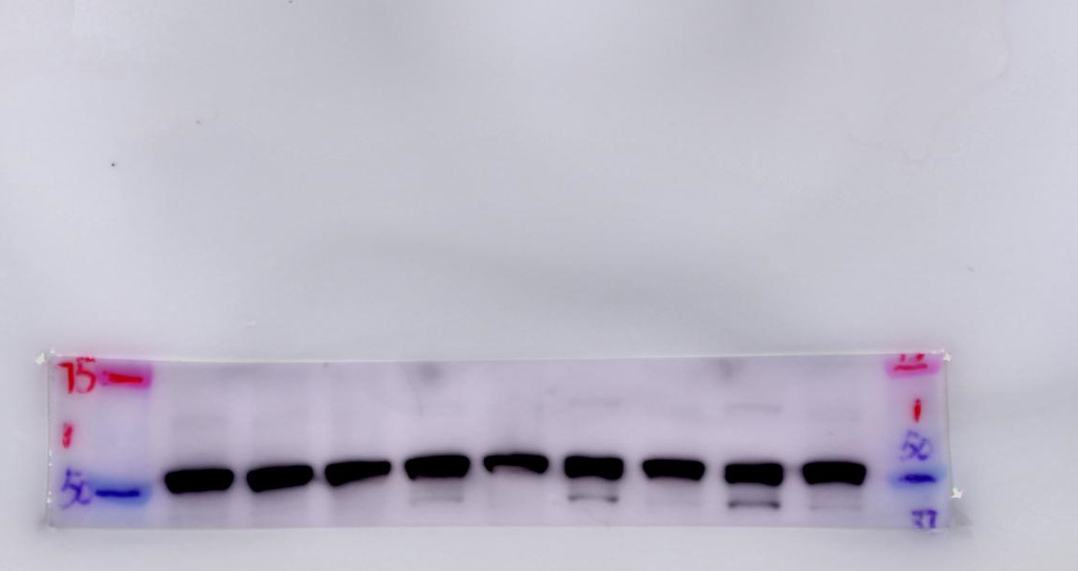

Figure	3C 11β-HSD1

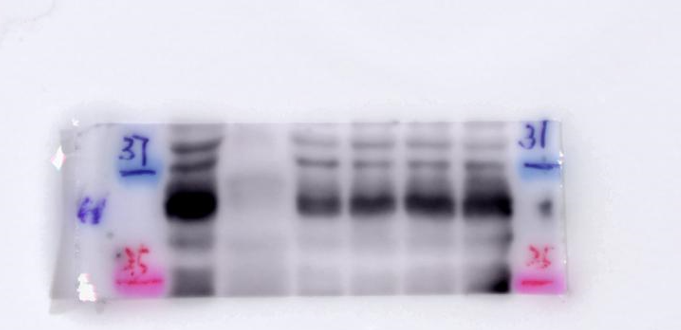

Figure	3C Tubulin

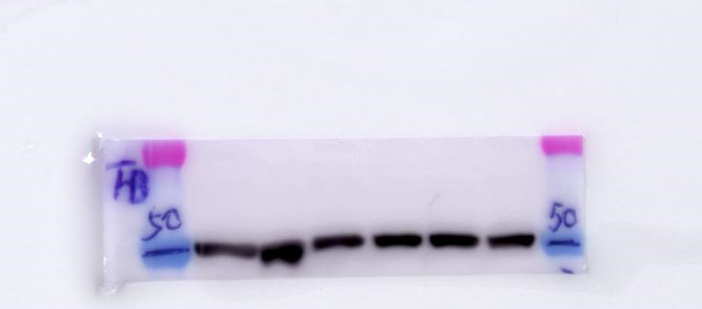

Figure	3D 11β-HSD1

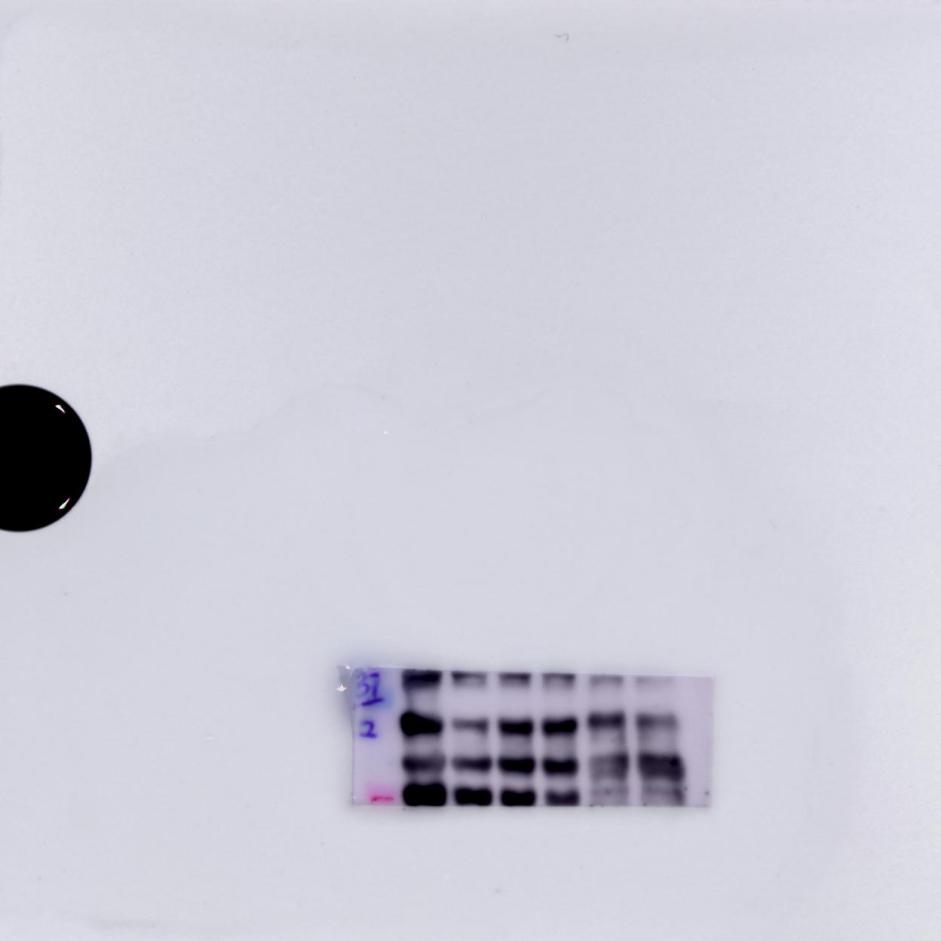

Figure	3D Tubulin

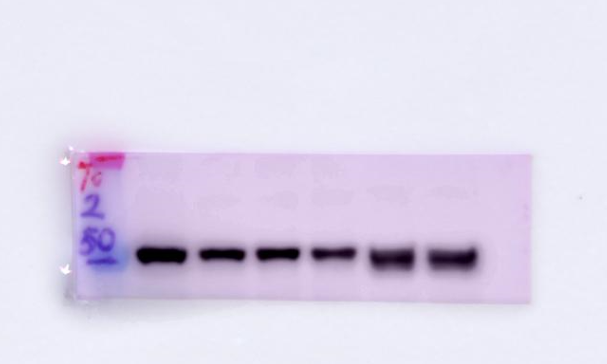

Figure	3E 11β-HSD1

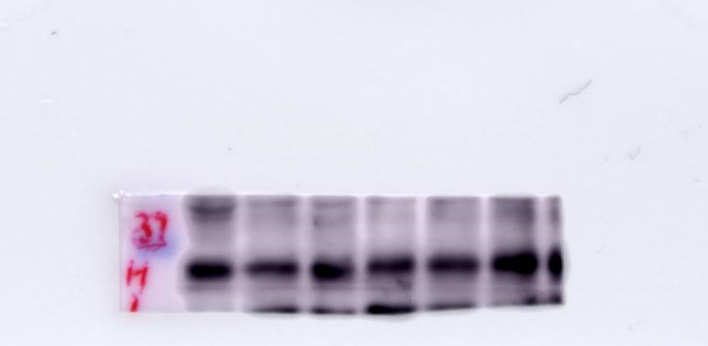

Figure	3E Tubulin

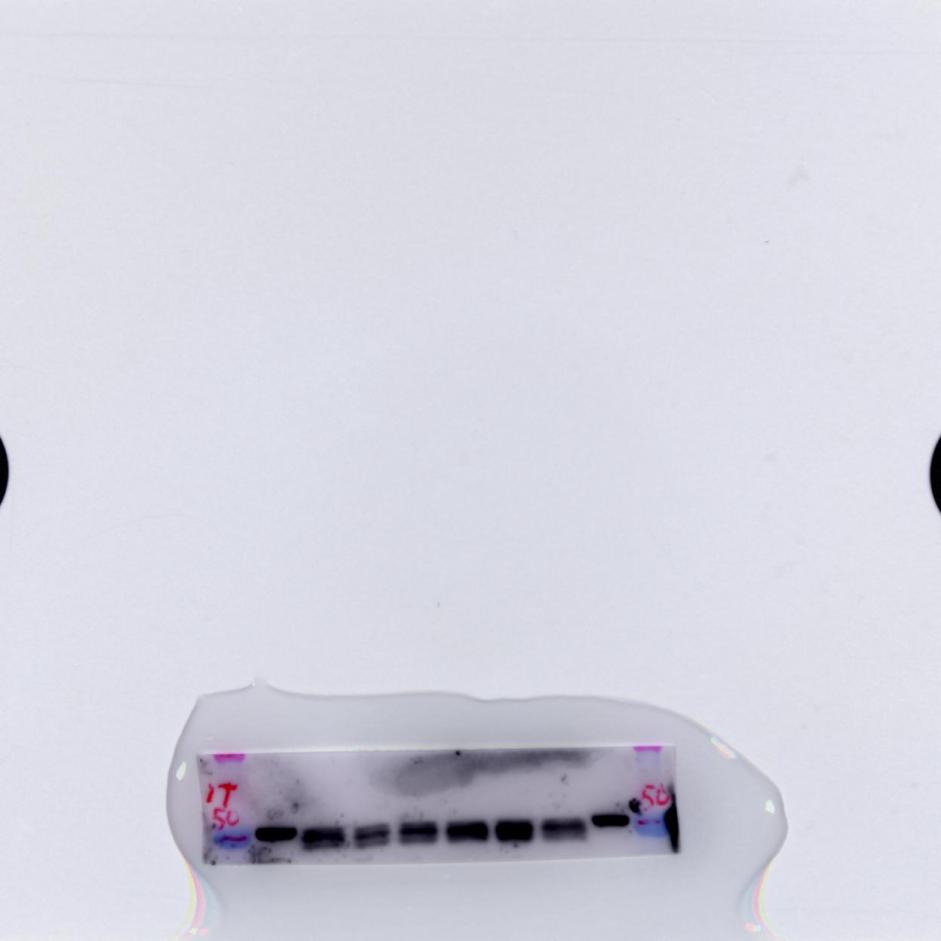

**Reference**

1	An, Z., Lv, W., Su, S., Wu, W. & Rao, Y. Developing potent PROTACs tools for selective degradation of HDAC6 protein. *Protein Cell* **10**, 606-609 (2019). <https://doi.org/10.1007/s13238-018-0602-z>

2	Lu, T. A comprehensive electron wavefunction analysis toolbox for chemists, Multiwfn. *J Chem Phys* **161** (2024). <https://doi.org/10.1063/5.0216272>

3	Lu, T. & Chen, F. Multiwfn: A multifunctional wavefunction analyzer. *Journal of Computational Chemistry* **33**, 580-592 (2011). <https://doi.org/10.1002/jcc.22885>

4	Valdes-Tresanco, M. S., Valdes-Tresanco, M. E., Valiente, P. A. & Moreno, E. gmx_MMPBSA: A New Tool to Perform End-State Free Energy Calculations with GROMACS. *J Chem Theory Comput* **17**, 6281-6291 (2021). <https://doi.org/10.1021/acs.jctc.1c00645>
